# The sugarless grape trait characterized by single berry phenotyping

**DOI:** 10.1101/2022.03.29.486323

**Authors:** Antoine Bigard, Charles Romieu, Hernán Ojeda, Laurent Torregrosa

## Abstract

For grape production, an important driver for the selection of varieties better adapted to climate fluctuations, especially warming, is the balance between fruit sugars and acidity. Since the past decades, temperature during ripening has constantly raised causing excessive sugars concentrations and insufficient acidity of the wine grapes in warmest regions. There is thus an increasing interest in breeding new cultivars, able to ripen at lower sugar concentration while preserving fruit acidity. However, the phenotyping of berry composition challenges both methodological and conceptual issues. Indeed, most authors predetermine either average harvest date, ripening duration, thermal time or even hexoses concentration threshold itself, to compare accessions at an hopefully similar ripe stage. Here, we have phenotyped the fruit development and composition of 6 genotypes, including 3 new disease-tolerant varieties known to produce wines with low alcoholic contents. The study was performed at single berry level from the end of green growth stage to the arrest of phloem unloading, when water and solute contents reach a maximum per berry. The results confirmed that sugarless genotypes achieve fruit ripening with 20-30% less hexoses than classical varieties, Grenache N and Merlot N, without impacting berry growth, total acidity or cations accumulation. Sugarless genotypes displayed a higher malic acid/tartaric acid balance than other genotypes with similar sucrose/H+ exchanges at the onset of ripening. Data suggest that sugarless phenotype results from a specific plasticity in the relationship between growth and the turgor imposed by organic acid accumulation and sugar loading. This opens interesting perspectives to understand the mechanism of grapevine berry growth and to breed varieties better coping with climate warming.

## Introduction

With an hexose concentration (glucose + fructose, [Hex]) higher than 1.1 mol.l^-1^ at ripe stage, grape is one of the richest fleshy fruits in sugars. [Hex] is accepted to depend on the GxExM interaction (Suter et al., 2021) and until recently, cultivars adaptation to local conditions was essentially reasoned according to the thermal time needed to reach specific vegetative and reproductive phenological stages, as budburst, flowering or fruit veraison (Parker et al., 2020). The selection of grapevine varieties is mainly driven by the balance between sugars and acidity (Torregrosa et al., 2017; Ollat et al., 2018; Duchêne et al., 2020). In cold climates, early ripening varieties are preferred to secure the accumulation of sugars and secondary metabolites before autumn. Conversely, in warm regions, late ripening varieties shifting fruit ripening to cooler days preserve organic acids (Rienth et al., 2016), anthocyanidin (Zhang et al., 2015) and aroma compounds (Alessandrini et al., 2018; Asproudi et al., 2016; Gutiérrez-Gamboa et al., 2018). But in practice a range of enological processes are implemented to correct sugars and/or acidity of the must, demonstrating that the supposed adaptation of the varieties based on thermal time phenology is oversimplified. Since the past decades, temperature during the period of grapevine fruit ripening has constantly increased, which may lead to excessive sugars and insufficient acidity in hot vine growing areas, such as mediterranean regions (Santillan et al., 2019, Bécart et al., 2022). Especially, with the strength and the rate of ongoing climate changes, it is critical to better objectify the development and the metabolism of the fruits to characterize their adaptation potential (Bigard et al., 2018 and 2020). There is thus increased interest in breeding new cultivars, able to ripen at lower sugar concentration while preserving fruit acidity.

Actually, warming doesn’t simply accelerate the whole ripening process, which would be easily solved by harvesting earlier but it decorrelates different aspects of ripening, accelerating malic acid breakdown (Rienth et al., 2016, Sweetmann et al., 2014) while inhibiting the accumulation of secondary metabolites such as anthocyanins, hence the decision to shift the date of harvest to higher [Hex] (Arrizabalaga et al., 2018). Moreover, comparing cultivars at the same developmental stage (the so-called ripe stage) raises both methodological and conceptual issues. In most comparative studies, authors predetermine an average harvest date, ripening duration, thermal time or even [Hex] threshold to compare accessions at an hopefully similar ripe stage, in contradiction with the recognized impact of GxE on these variables (Liu et al., 2007; Dai et al., 2011; Duchêne et al., 2012). To circumvent this inconsistency and lack of consensus, the moment at which berry phloem unloading stops was recently proposed as a relevant definition of ripe stage both on the physiological and transcriptomic point of views (Bigard et al., 2018 and 2020; Shahood et al., 2015 and 2020; Savoi et al., 2021). This key transition which marks the arrest of the most intensive fruit biochemical pathways is associated with the transcriptional extinction of many genes encoding, amongst other, sugar transporters, aquaporins and cell wall-related enzymes (Savoi et al., 2021). Unfortunately, this developmental stage can’t be directly inferred from [Hex] kinetics, which continue to evolve after the completion of sugar storage (or accumulation) due to subsequent berry shriveling. Thus, additional information on berry growth is required to address the net rates of sugar accumulation and malic acid breakdown in berries, together with their respective timings. Very recently, single berry phenotyping approaches improved our understanding of berry growth and metabolism during ripening, avoiding the biases due to berry asynchronicity (Shahood et al., 2020; Savoi et al., 2021). According to this new paradigm, this study aims to decipher the genetic differences existing for [Hex] and fruit acidity in a set of genotypes encompassing traditional varieties and new hybrids exhibiting low [Hex] at harvest (Escudier et al., 2017; Ojeda et al., 2017).

## Materials and method

### Plant Material and sampling method

Experiments were performed outdoors at the INRAE Pech Rouge experimental unit (Gruissan, France, 43.14’/3.14’’W) under a semi-arid Mediterranean climate in 2017 (temperature, rainfall and evapotranspiration data described in Alem et al., 2021). Experimental plots were managed through drip irrigation keeping the predawn leaf water potential (ΨPD) higher than −0.5 MPa (Giorgi & Lionello, 2008) to correspond to a moderate water stress. The set of the genotypes encompassed 3 traditional varieties Grenache N, Merlot N, and Morrastel N (https://plantgrape.plantnet-project.org/fr/) and 3 new disease-resistant varieties deriving from 4 (3197-81B, 3197-373N) or 5 (3184-1-9N) backcrosses of *Muscadinia rotundifolia* with *V. vinifera* varieties (Bouquet al., 1980). These 3 last genotypes are known to display a reduced [Hex] at harvest allowing the production of wines at low ethanol levels, called VDQA, “***V****ins* ***D****e* ***Q****ualité à teneur modérée en* ***A****lcool*” (Ojeda et al. 2017). In the rest of the manuscript, the names G5, G7 and G14 will be respectively used for 3197-81B, 3197-373N and 3184-1-9N. From 2 weeks before the first signs of berry softening to 2 weeks later and during the rest of the ripening period over 1 week after fruit shriveling, respectively 60 and 30 berries were weekly and randomly collected. Whole berries were sampled between 9 and 11 AM by cutting the fruit peduncle just below the calyx, maintained in a plastic bag in a cool place and analysed in the same day.

### Berry firmness and composition

Firmness was monitored with a digital penetrometer called Pénélaup™ (Abbal et al., 1992; Robin et al., 1997) as described in Shahood et al. (2020). Immediately after firmness measurement, berries were immersed in 4 times their weight in 0.25N HCl. Seeds were immediately removed and samples incubated 48h. Samples were vigorously shaken and a first 100 *μ*L aliquot was 11 times diluted with 8.3 10^−3^ N acetic acid (internal control) + 16.4 10^−3^ N sulphuric acid and centrifuged 5 min’ at 18,500 g at 20°C, supernatants were injected for HPLC to quantitate glucose, fructose, malic and tartaric acids through a Biorad aminex-HPX87H column also described in Shahood et al. (2020). A second 100 *μ*L aliquot was diluted 10 times in water and then 3 min centrifuged at 12000 rpm (20°C). Ten μl clear supernatant was then injected in the HPLC through a Waters^®^ IC-Pak Cation M/D 3.9×150 mm column with same parameters used in Bigard et al. (2020) in order to obtain Potassium ([K^+^]), Magnesium ([Mg^2+^]) and Calcium ([Ca^2+^]) concentrations. Titratable acidity (TA) was calculated as the sum of malic and tartaric acids minus K^+^ in mEq.L^-1^.

### Data normalisation and presentations

R^®^ software (version 4.1.2) was used to build graphical representations and to analyse the data (R Core Team, 2017). Main packages used for this study were “ggplot2” (Version 3.3.5), “car” (Version 3.0-12) and “rcompanion” (Version 2.4.6).

## Results and discussion

The grapevine displays small fruits clustered in grapes presenting a huge internal asynchrony (Gouthu et al., 2014; Doumouya et al., 2014; Bigard et al., 2019; Shahood et al., 2020), as illustrated in the next figures. Extrapolating single fruit metabolic traits from the averages values observed on the population of berries led to biased kinetic interpretations and chimeric metabolic concepts. In the grapevine where the berry is the only truly relevant physiological unit, the most accurate method would be to non-destructively characterize each single fruit kinetics (Castellarin et al., 2015; Savoi et al., 2021), which is only possible for morphological attributes (fruit color and volume) and firmness. Regarding solute accumulation, hypodermal sampling led to excessively elevated fluxes possibly resulting from injuring the berries, so its validity was questioned (Coombe, 1992). Destructive sampling of density sorted berries or large sets of individual fruits is then required to get accurate physiological insights on grapevine fruit development (Rolle et al., 2011; Bigard et al., 2019; Shahood et al., 2020). Here, we phenotyped the ripening of 3 new sugarless genotypes (G5, G7 and G14) and 3 traditional varieties (Grenache N, Merlot N, Morrastel N) though the destructive chemical analysis of thousands of single berries as in Shahood et al. (2020). Since it was not possible to measure each individual flowering or softening dates, data are interpreted as a function of berry sugar concentration, a proxy for internal time of fruit ripening (Rienth et al., 2016).

### Berry development and sugar accumulation

**Figure 1** shows the evolution of berry weight and tartaric acid concentration during ripening of Morrastel N, representative of the panel of varieties described in this study. Observed trends are typical of *V. vinifera* genotypes with a nearly doubling berry volume and a twofold dilution of tartaric acid during ripening. As described in Rienth et al. (2016) and in Bigard et al. (2020), the dilution of tartaric acid appears to be a relevant indicator of berry relative growth as its quantity doesn’t evolve during and after ripening (Ruffner, 1982; Lang & Thorpe, 1989; Terrier & Romieu, 2001; Rienth et al., 2014; Rösti et al., 2018; Burbidge et al., 2021). When the uploading of sugars and water in the berries stops (Coombe & McCarthy, 2000; Conde et al., 2007; Savoi et al., 2021), hexoses and tartaric acids just continue to concentrate due to evaporation. This results in a linear regression passing through the origin as in **figure 1B**. Tartaric acid concentration was obviously less heterogene than fresh berry weight facilitating the identification of the max berry volume stage. With this method we could determine the berry [Hex] at the maximum fruit volume for each variety, i.e. : 920 mmol.L^-1^ for G5, 900 mmol.L^-1^ for G7, 800 mmol.L^-1^ for G14, 1125 mmol.L^-1^ for Grenache N, 1125mmol.L^-1^ for Merlot N and 1000 mmol.L^-1^ for Morrastel N. We used the 10 closest berries for each genotype and both stages (end of green growth and max berry volume) to perform statistics as recorded in **Table 1**. For the end of the green growth period, the 10 berries showing the highest malic acid concentration were selected.

**Table 1.**
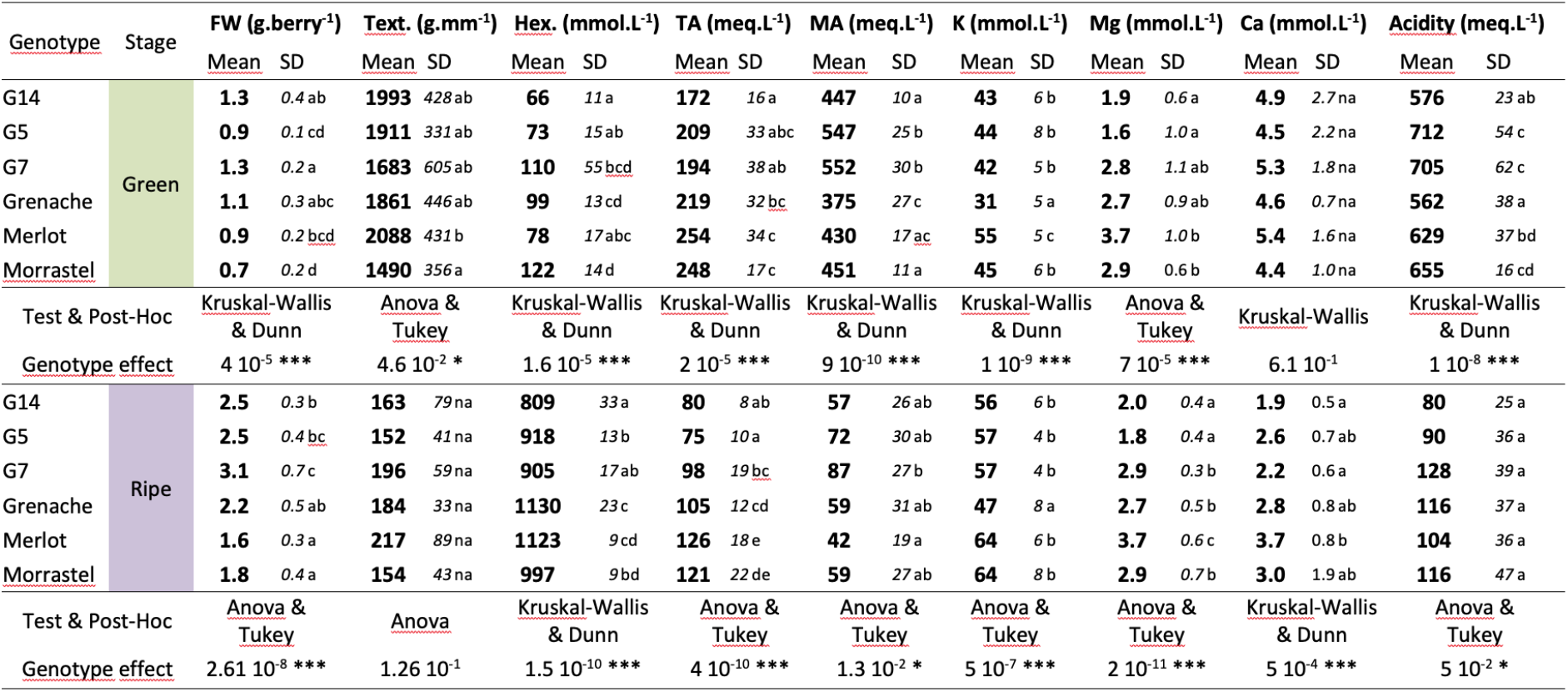
Berry weight, firmness and composition of the 6 studied genotypes at the end of green growth phase and at physiological ripe stage. FW (Fresh weight), Text. (Texture), Hex (Hexoses), TA (Tartaric acid), MA (Malic acid). Statistical significance: * (p<0.01), ** (p<0.001), *** (P<0.0001), na (p>0.01).

**Fig. 1.**
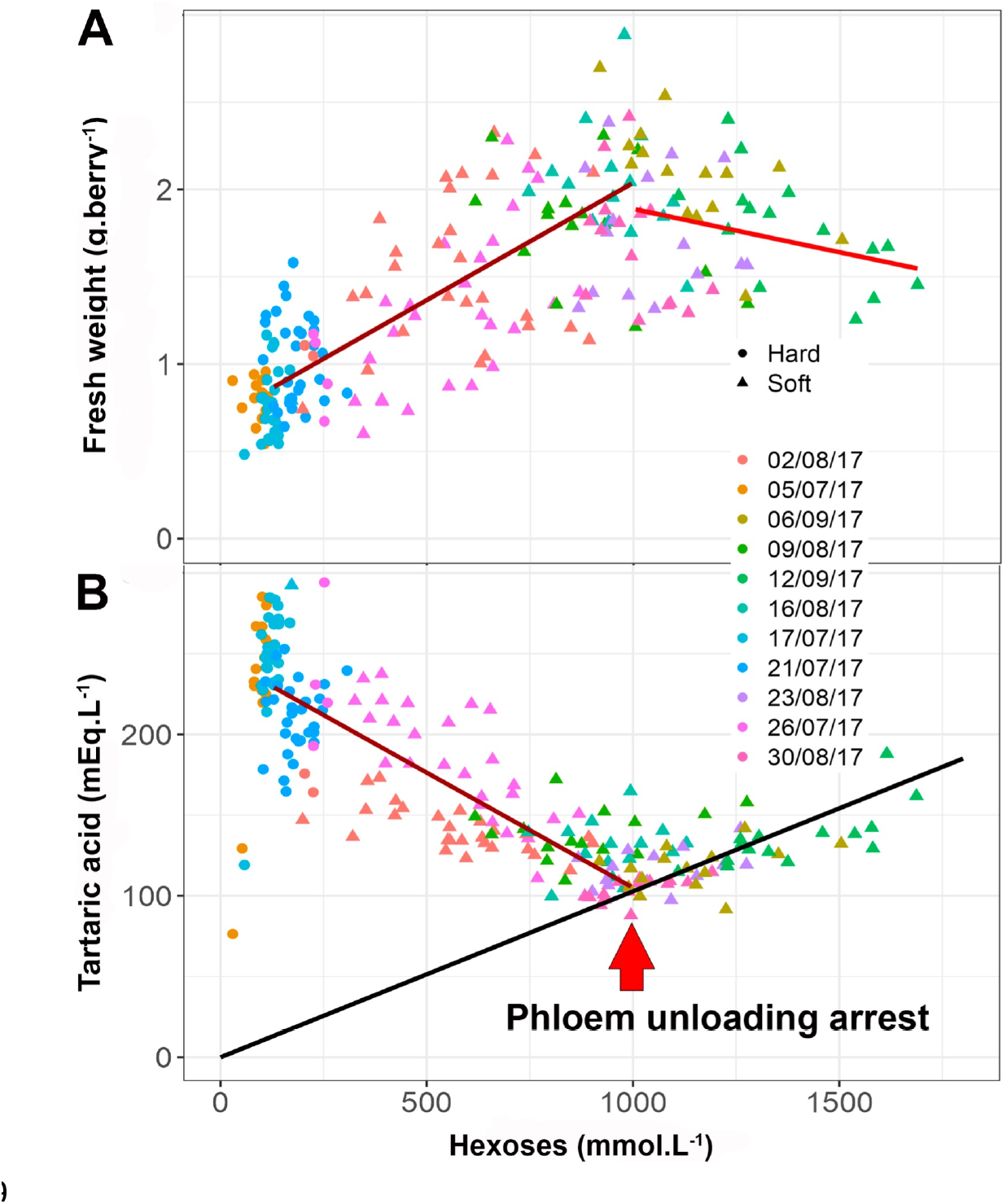
Evolution of fresh weight (A) and tartaric acid concentration (B) of single berries of Morrastel during ripening.

Despite genotypic differences in berry weight at green stage (**Fig. 2A**), the 6 genotypes displayed the classical fruit expansion kinetics during ripening (Coombe, 1976). The relative volume increment (i.e. Vripe/Vveraison) was obtained using all combinations possible between green and ripe selected berries and ranged from 1.8 ^+^/- 0.5 to 3.1 ^+^/- 0.7, for Merlot N and G5 respectively. Statistical analyses revealed that G5 had the highest berry growth during ripening followed by G7 and Morrastel N, then G14 and Grenache N with Merlot N has the least.

**FIg. 2.**
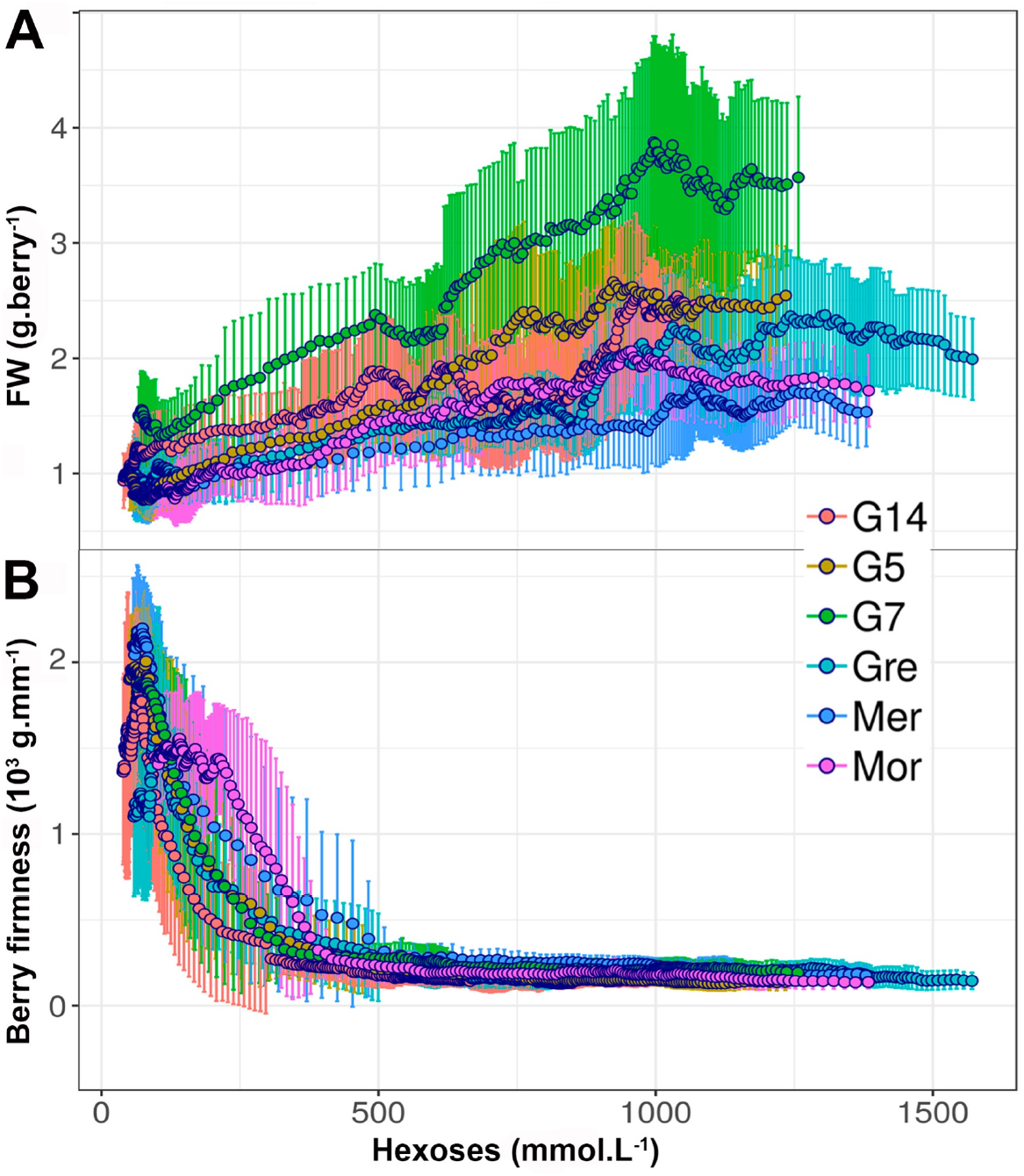
Evolution of the berry fresh mass (A) and firmness (B) during the fruit ripening of 6 grapevine varieties.

Whatever the genotype, we observed a considerable heterogeneity in berry size at a similar [Hex], as previously reported (Gouthu et al. 2014; Vondras et al., 2016; Bigard et al. 2019; Shahood et al., 2020). Both the maximum volume of the berries and [Hex] at this stage varied according to the variety (**Fig. 2A**). [Hex] increased during ripening, from values around 100 mmol.L^-1^ at the end of the green growth phase to 0.8 to 1.1 mol.L^-1^ at physiological ripe stage (**Table 1**). During berry ripening, with ca 0.5 M each, glucose and fructose became the major osmoticums as reported before in *V. vinifera* (Hawker et al., 1976; Liu et al., 2006; Shiraishi et al., 2010). [Hex] observed in this study at phloem arrest for the Merlot N and Grenache N are lower than the usual concentration threshold of 1.2 to 1.5 mol.L^-1^ [Hex] at which the industry considers the berries as technologically ripe. Kliewer (1967) reported a range from 1 mol.L^-1^ to 1.5 mol.L^-1^ [Hex] as the technical ripe grape for usual *V. vinifera* varieties. This apparent discrepancy is due to the very common practice to push grapes towards over-ripeness to get more redfull, aromatic wines and concentrated wines (Antalick et al., 2021). In the absence of supplementary physiological landmarks, the use of [Hex] for comparative studies is very hazardous, as this parameter steadily increases after phloem unloading arrest because of fruit shriveling (Friend et al., 2009; Shahood et al., 2020; **Fig. 2A**).

Sucrose unloading in berries of all genotypes dramatically increased at softening, or relaxation of turgor pressure (**Fig. S1**). All genotypes displayed a glucose/fructose higher than 2.2 before fruit softening, then the ratio progressively converged to 1 as reported in other *V. vinifera* varieties (Varandas et al., 2004). No specific metabolic trends could be observed in the G genotypes. It is known that berry glucose/fructose balance which depends on grapevine organs and developmental stage can be used as a metabolic indicator of fruit ripening (Kliewer et al., 1966). During green growth, the preferential use of fructose is obvious, leading to elevated G/F ratio. At softening, the import of sucrose dramatically accelerates, exceeding metabolic needs, insofar as malic acid replaces sugar as a respiratory substrate. Consequently, the G/F ratio rapidly tends to 1 (Amerine and Thoukis, 1958; Liang et al., 2011; Houel et al., 2015).

Within a range of extreme varieties and offsprings Bigard et al. (2018) showed that [Hex] can vary from 750 to 1350mmol.L^-1^ when solute unloading stops just before berry shriveling. Here, considering phloem arrest as the physiological ripe stage, G genotypes displayed [Hex] levels between 0.8 and 0.9 mol.L^-1^, Morrastel N was at 1 mol.L^-1^, Merlot N and Grenache N showed the highest [Hex] (> 1.1 mol.L^-1^). These data agree with previous results obtained at the whole berry population levels with similar genotypes (Ojeda et al., 2017; Bigard et al., 2019). Interestingly, the final quantity of sugar per fruit unit in sugarless genotypes is of the same magnitude as classical varieties, i.e : 2.8 +/- 0.3 mM (G7), 2.5 +/- 0.6 mM (G5), 2.3 +/- 0.4 mM (G5), 2.0 +/- 0.3 mM (G14), 1.8 +/- 0.3 mM (Merlot) and 1.8 +/- 0.4 mM (Morrastel) per berry. Before softening, berry firmness showed some differences in berries at the end of the green growth phase with the Morrastel N and the Merlot N respectively showing the least and the most firm fruits (**Table 1**). From the green growth phase, mechanical properties of the berries evolved in the same way for all genotypes (**Fig. 2B**). Berries soften rapidly at the beginning of ripening to reach a low level of firmness before 500 mmol.L^-1^ of [Hex], i.e. more or less mid ripening. Then, with a slow and continuous decrease of the firmness up to physiological ripe stage and over-ripening, no firmness differences could be observed between sugarless genotypes and traditional cultivars (**Table 1, Fig. S1**). Therefore, although firmness is widely accepted as a sensitive and early indicator of the onset of ripening (Coombe et al. 1992; Abbal et al., 1992; Castellarin et al., 2015; Shahood et al., 2020; Bigard et al., 2020), this parameter can’t be used to tag the transition at phloem unloading arrest. Consequently, the only way to non destructively determine the shift from fruit expansion to shriveling remains the monitoring of berry growth. As discussed below, this can be done indirectly, using Tartrate concentration

**Fig. S1.**
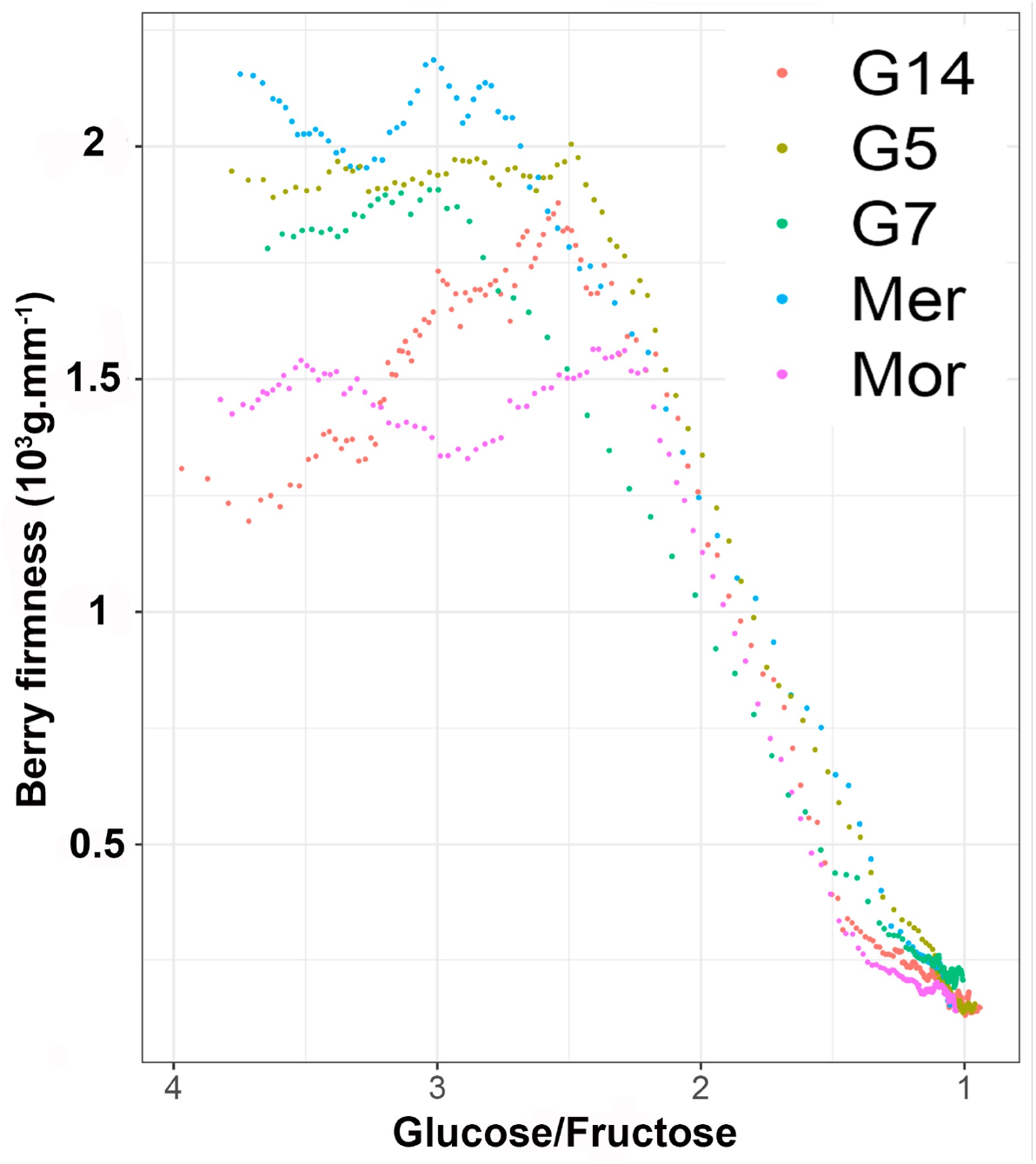
Evolution of the berry firmness depending on glucose/fructose ratio during the fruit ripening.

### Evolution of the main determinants of fruit acidity

#### Tartaric acid

During ripening, tartaric concentration depended on the variety and the stage of berry development (**Table 1**). At the onset of ripening, tartaric acid concentration was 25% lower (150 vs 200 mEq.L^-1^) in all G genotypes. Tartaric acid dilution (**Fig. S2**) proceeded at negligible rate before 220 mmol.L^-1^ (G5, G14) to 300 mmol.L^-1^ [Hex] (Grenache N) and then accelerated confirming the delay between berry softening and growth resumption (Castellarin et al., 2015; Shahood et al., 2020 and other literature cited in this paper).

The change in tartaric acid from the starting to the end of ripening (**Fig. 3**), is consistent with a first two to three fold dilution, followed by concentration due to shriveling, leading to a linear increase in sugar and tartaric acid passing the origin. At the arrest of phloem unloading, the concentration of tartaric acid reached a minimum ranging from 75 ^+^/- 10 mEq.L^-1^ (G5) to 126 ^+^/- 18 mEq.L^-1^ (Merlot N). Grenache N displayed a 104 ^+^/- 12 mEq.L^-1^ concentration in tartaric acid at the ripe stage.

**Fig. 3.**
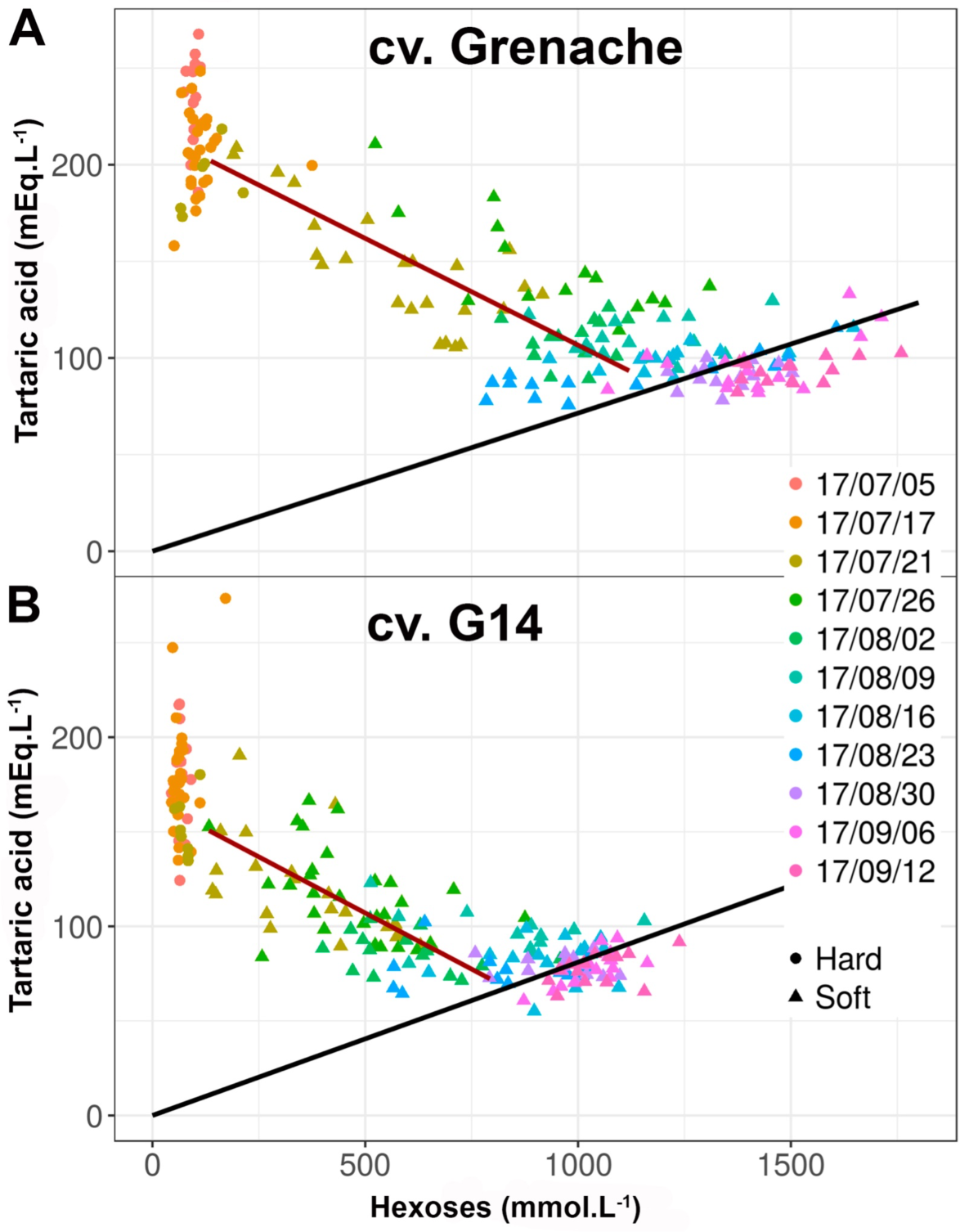
Evolution of the tartaric acid concentration during berry ripening of Grenache (A) and G14 (B). Lines corresponding to linear fitting during and after phloem unloading.

**Fig. S2.**
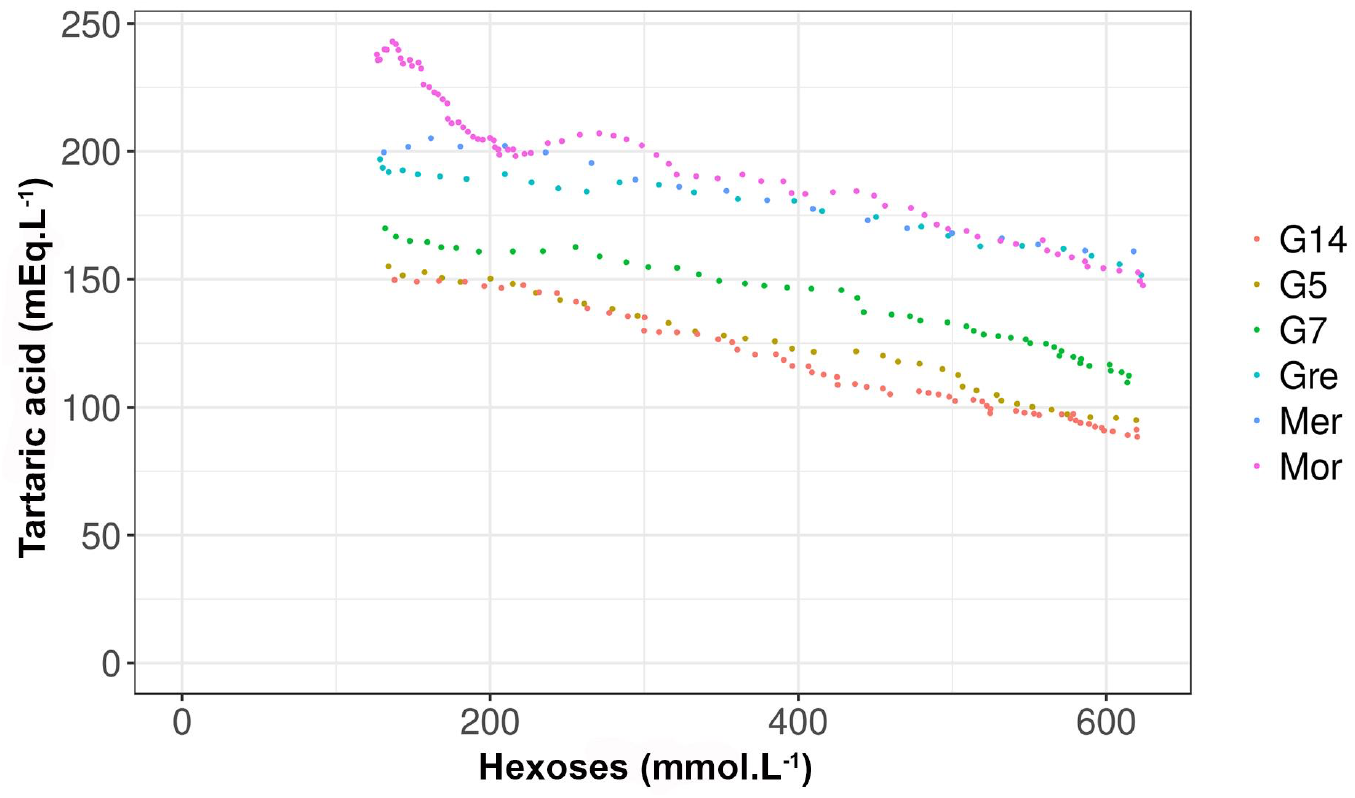
Evolution of the tartaric acid concentration at the early stages ([Hex] from 125 and 625 mmol.L^-1^) of fruit ripening of 6 grapevine varieties.

Tartaric acid is the first organic acid to be accumulated during young berry development and remains one of the main acids in ripe fruits of *Vitis vinifera* (Amerine et al., 1965; Kliewer, 1966). In this study performed through single berry analysis, tartaric acid ranged from 170-250 mEq.L^-1^ at the beginning of ripening to decrease to 70-120 mEq.L^-1^ at phloem stop as previously observed (Bigard et al., 2019). Statistical analyses showed that, at the onset of ripening, sugarless genotypes already display a lower tartaric acid concentration than traditional cultivars. This trend is amplified at maximum volume stage due to the highest expansion and resulting dilution (**Table 1**). Morrastel N, also called Graciano in Spain, is a traditional cultivar producing wines rich in polyphenols with moderate ethanol levels (Ramos and Martinez de Toda, 2021). We confirm here, through this study performed at single berry level, that Morrastel N can produce ripe fruit at lower [Hex], i.e. below 1 mol.L^-1^, in comparison to other traditional varieties.

#### Malic acid

Malic acid concentration peaked at 370-550 mEq.L^-1^ at the very end of green growth period and then decreased to less than 90 mEq.L^-1^ at maximum berry volume whatever the genotype (**Table 1**). At the onset of ripening, conversely to tartaric acid, malic acid concentration was higher in sugarless genotypes than in traditional varieties. At the arrest of phloem unloading, the concentrations in malic acid ranged from 42 ^+^/-20 mEq.L^-1^ (Grenache N) to 87 ^+^/- 26 mEq.L^-1^ (G7), with no obvious genotypic effects. After phloem arrest, conversely to tartaric acid, malic acid concentration stayed stable or even slightly decreased (**Fig. 4**), as observed in previous studies performed at berry population levels (Ojeda et al., 2017; Bigard et al., 2019).

**FIg. 4.**
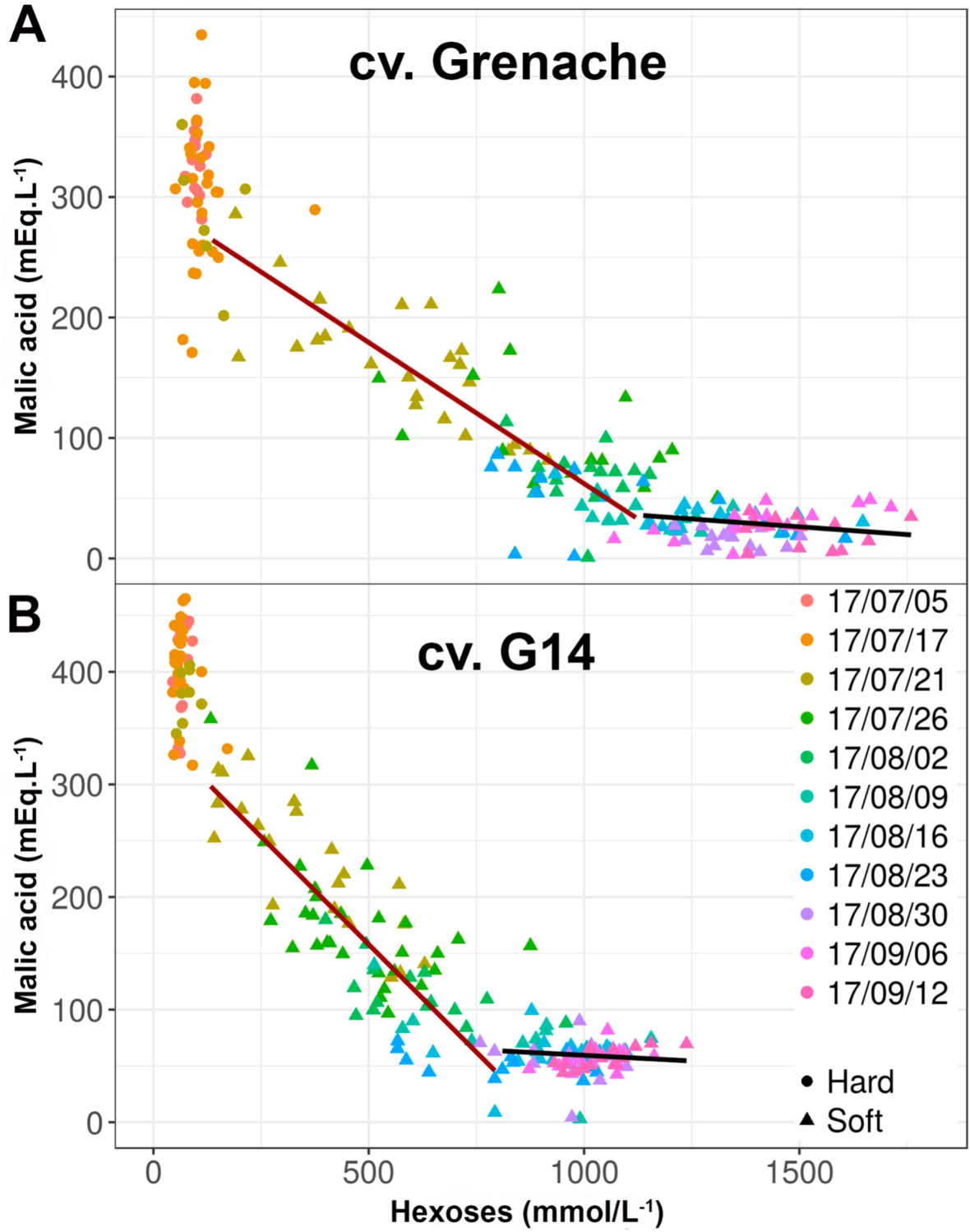
Evolution of the malic acid concentration during berry ripening of Grenache (A) and G14 (B). Lines corresponding to linear fitting during and after phloem unloading.

Malic acid respiration provides a major fraction of the energy during early ripening (Famiani et al., 2014; Shahood, 2017), hence a faster decrease in concentration than if only dependent on dilution due to berry growth. Here, despite different starting points (**Table 1, Fig. S3)**, the decrease of malic acid during the first phase of ripening (i.e. from 250 to 800 μmol.berry^-1^ [Hex]), was characterized by an initial slope of −1 mEq per 2 hexoses, noticeably similar in the 6 genotypes (**Fig. S3**). During early ripening, the initial changes in the respective amounts of malic acid and sugar per fruit (concentration x volume; **Fig. 5**) are fully consistent with the activation of a sucrose/H^+^ exchanger on the tonoplast of all *V. vinifera* cultivars investigated, including sugarless genotypes, which generalizes our quantitative data on Syrah and Pinot (Shahood et al., 2020). In single berries, the corresponding genes are strongly expressed until phloem arrest (Savoi et al., 2021). At the beginning of ripening, the sucrose/H+ exchange is electro-neutralized by the release of vacuolar malate, as detected here, while more and more H+ must be redirected to the vacuole as malic acid vanishes, as illustrated by the progressive activation of vacuolar ATPase and PPiase (Terrier et al., 2001).

**Fig. S3.**
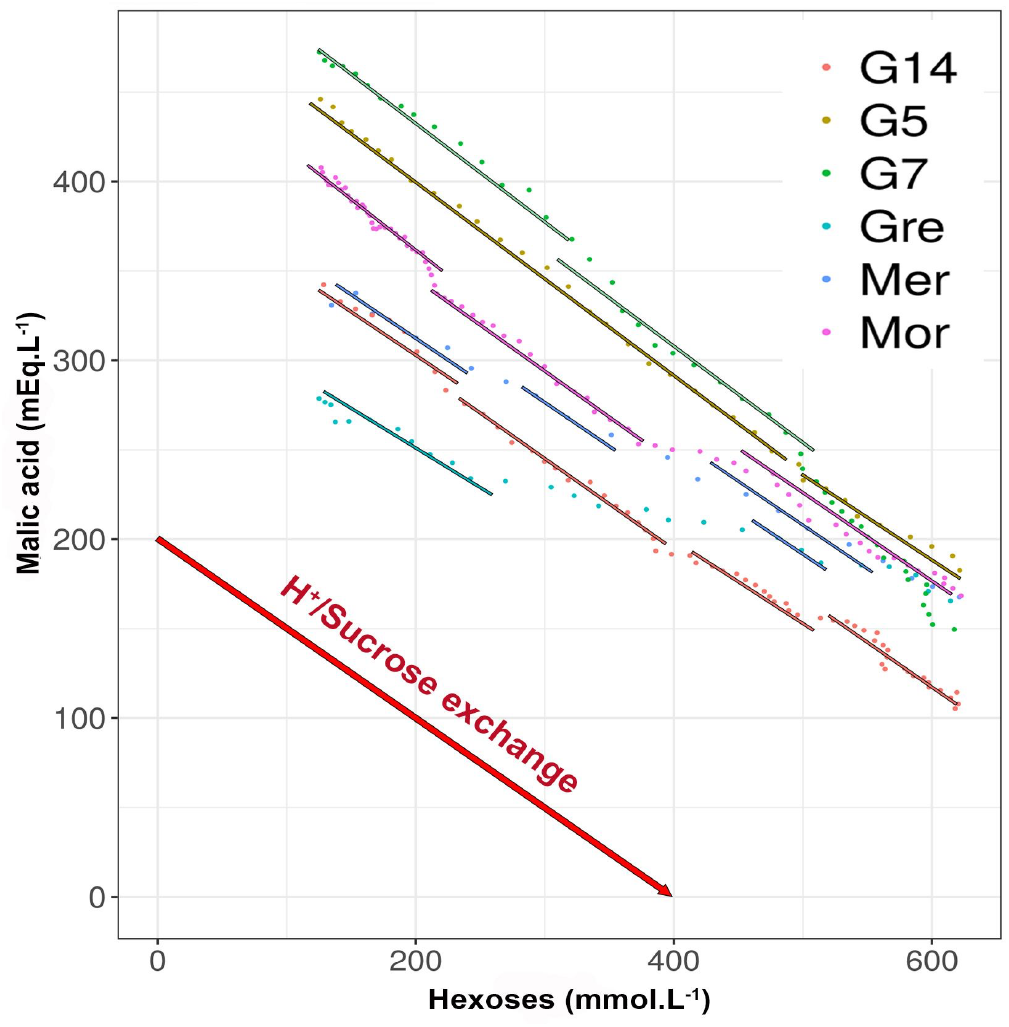
Repeatability of the malic acid/sugar exchange during early berry ripening (125-625 mmol.L-1 [Hex]) of 6 grapevine varieties.

**Fig. 5.**
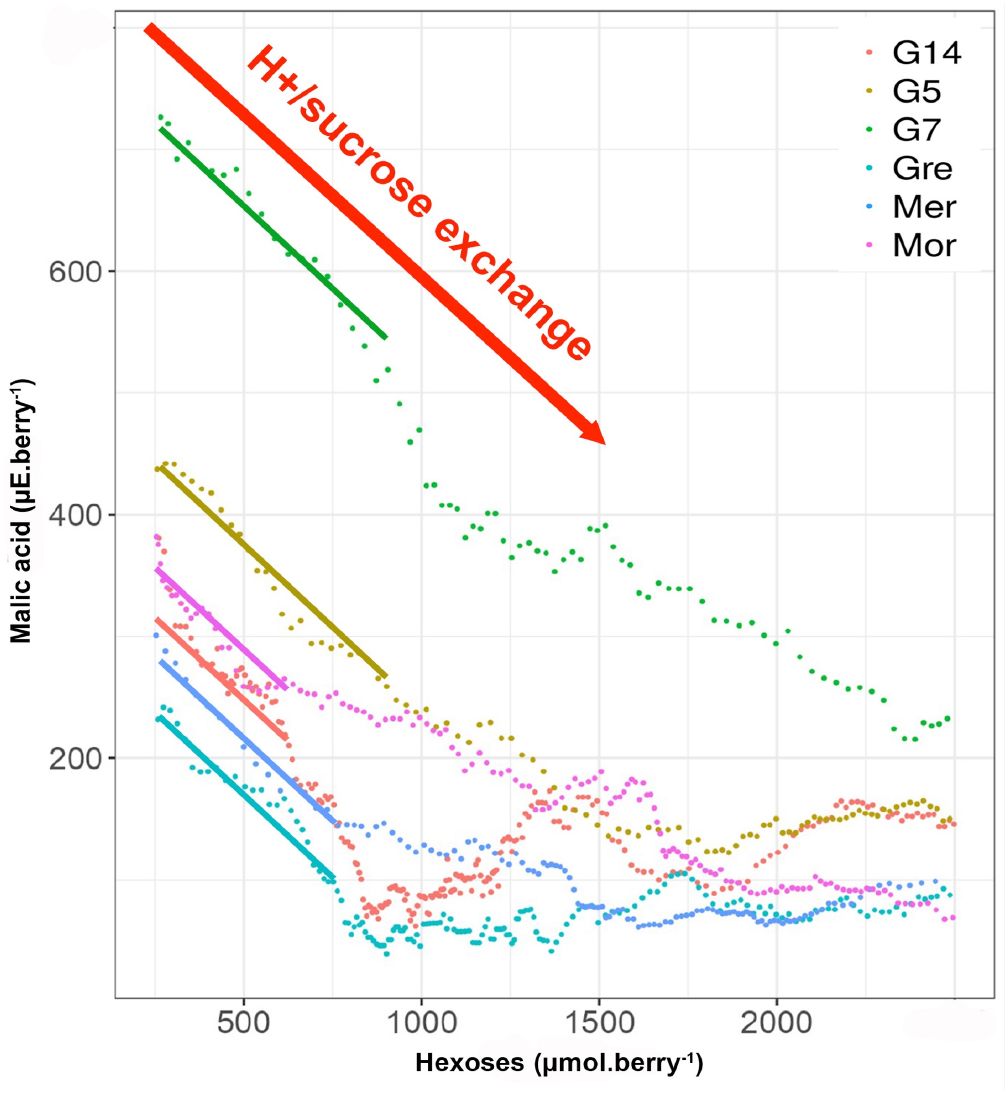
Repeatability of the malic acid/sugar exchange expressed in quantity per fruit during berry ripening of 6 grapevine varieties.

When the cumulative evolution of tartaric + malic acids was monitored during early ripening (**Fig. S4 A**), no specific behaviors could be observed in the sugarless genotypes in comparison to the other varieties. Considering that, according to their concentrations, sugars and organic acids are the main contributors to the osmotic potential of the berry (Matthews et al., 1987), present results totally exclude that the reduction in sugar concentration may be compensated by a greater accumulation of organic acids in the sugarless genotypes. As mentioned before, sugarless genotypes displayed lower tartaric and higher malic acid than traditional controls and consequently an higher malic acid/tartaric acid ratio (**Fig. S4 B**).

**Fig. S4.**
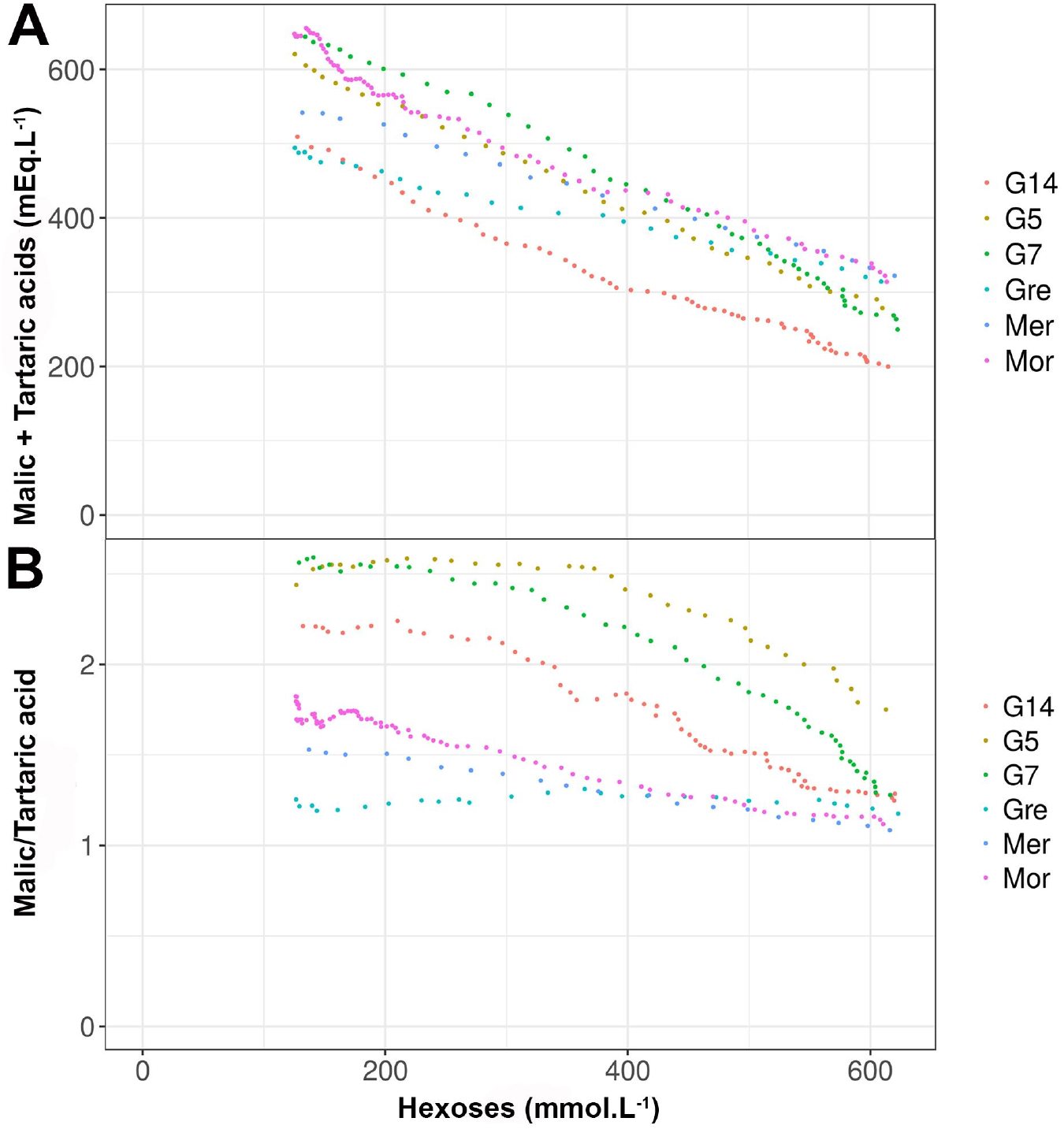
Evolution of the sum of malic and tartaric acid concentration (A) and malate/tartrate (B) during early berry ripening (125-625 mmol.L^-1^ [Hex]) of 6 grapevine varieties.

#### Potassium

K^+^, the main cation in the grapevine fruit, is accumulated during both phases of growth (Cuellar et al., 2013). Here, concentrations before ripening ranged from 31 ^+^/- 5 mEq.L^-1^ for Grenache N to 55 ^+^/- 5 mEq.L^-1^ for the Merlot N (**Table 1**). During ripening, [K^+^] increased moderately during ripening (**Fig. S5**) with increments ranging from 16% (Merlot N) to 50% (Grenache N) both genotypes displaying the lower and the higher levels of [K^+^] at the ripe stage, respectively 47 ^+^/- 8 mEq.L^-1^ for Grenache N and 64 ^+^/- 6 mEq.L^-1^ for Merlot N. After the period of phloem loading arrest, as for tartaric acid, [K^+^] steadily increased in link to water loss associated with shriveling (**Fig. 6**).

**FIg. 6.**
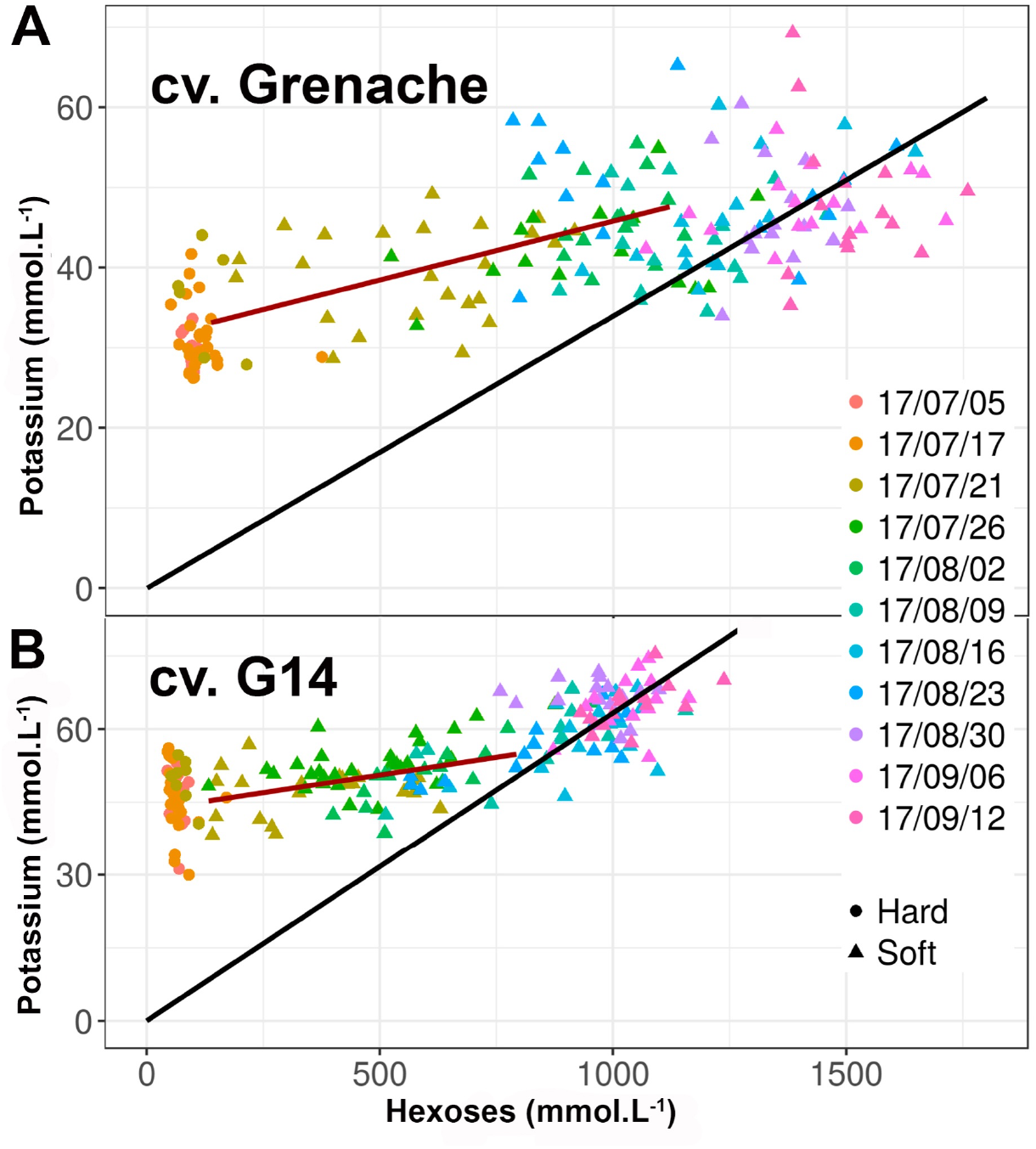
Evolution of the potassium concentration during berry ripening of Grenache (A) and G14 (B). Lines corresponding to linear fitting during and after phloem unloading.

**Fig. S5.**
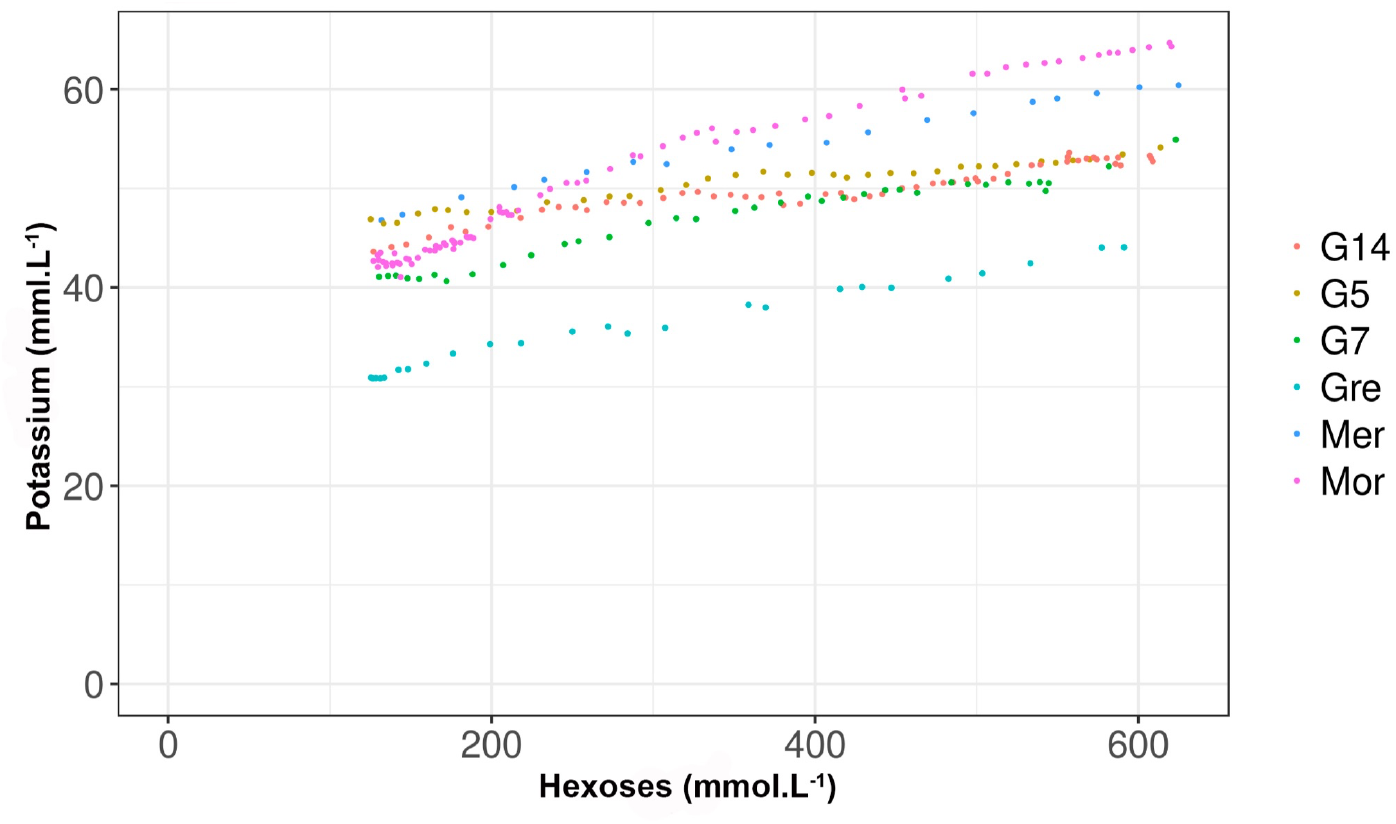
Evolution of the potassium concentration during early ripening (125-625 mmol.L^-1^ [Hex]) of 6 grapevine varieties.

After main organic compounds, K^+^ is the fourth contributor to berry osmotic potential, also neutralizing a fraction of organic acids (Storey, 1987; Rogiers et al., 2017). During early stages of development, low sugar accumulators (Morrastel N included) had a higher [K^+^] than high accumulators (**Table 1**). This element being mainly accumulated in the skin may play a role in the difference in elasticity from G genotypes. At the end of phloem unloading the average [K^+^] is around 50 mmol.L^-1^ in the 6 genotypes, suggesting a strong homeostasis for this element. During the phloem unloading period from veraison to max berry volume, as observed at population level with other genotypes (Bigard et al., 2020), K^+^ concentration increased 20-40 times less than hexoses (**Fig. 5, Fig. S5, Table 1**).

This obviously contradicts the so-called “massive” K^+^ import in the ripening berry (Villette et al., 2020). As recently discussed by Savoi et al. (2021), the belief that K^+^ transport would compensate for an intrinsic deficiency in the energisation of sugar imports is not supported by experimental data. In this respect, the simultaneous and parallel increases in [Hex] and [K^+^] observed after the arrest of phloem unloading, isn’t indicative of a co-transport mechanism, but only results from a net water loss and berry shriveling. Despite significant progress in the understanding of the import of potassium in grapevine berries (Rogiers et al., 2017; Villete et al., 2020; Savoi et al., 2021), the putative mechanistic links between potassium and sugar imports still remain speculative.

#### Evolution of the fruit acidity

Green berries displayed a high acidity **(Fig. 7, Table 1)** ranging from 560 ^+^/- 40 mEq.L^-1^ (Grenache N) to 710 ^+^/- 50 mEq.L^-1^ (G5). As the results of tartaric acid dilution, malic acid respiration and dilution, and slight K^+^ accumulation, acidity was reduced to 80 ^+^/- 25 mEq.L^-1^ (G14) to 130 ^+^/- 40 (G7) at the ripe stage with no statistical differences between genotypes. Noticeably, the total acidity tended to increase very late the ripe stage for all genotypes, ca 1250 mmol.L^-1^ [Hex] in Grenache N, and 1000 mmol.L^-1^ [Hex] in G14.

**FIg. 7.**
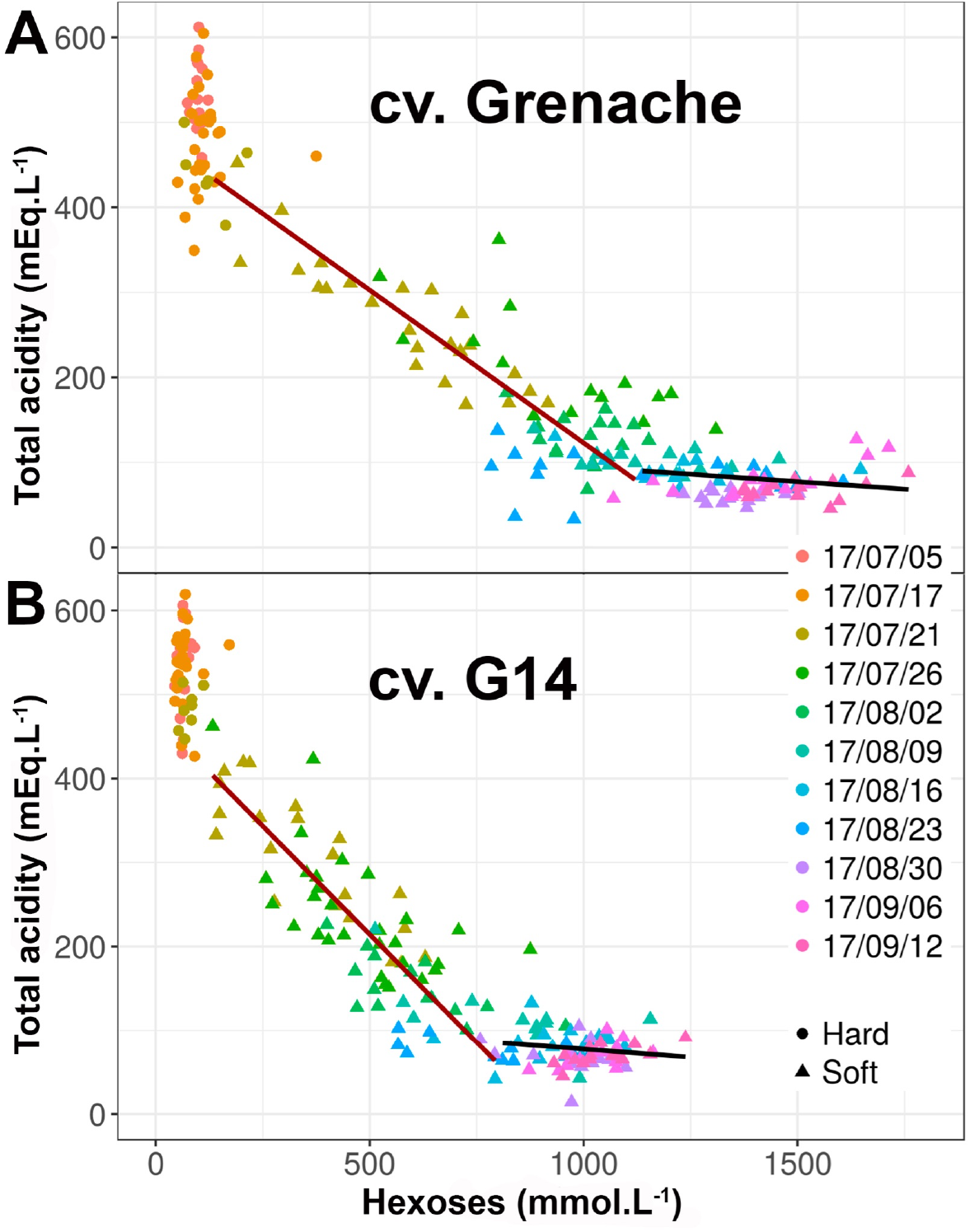
Evolution of the juice total acidity during berry ripening of Grenache (A) and G14 (B). Lines corresponding to linear fitting during and after phloem unloading.

Acidity is a major challenge for wine quality (Champagnol, 1984; Sweetman et al., 2014; Ollat et al., 2018). The effect of temperature on grape acidity is well documented (Kliewer and Lider, 1970; Butrose et al., 1971; Seguin et al., 2004; Rienth et al., 2016). By virtue of the electroneutrality principle, titratable (or total) acidity represents the difference between acids (mainly tartaric and malic in grapevine) and cations (mainly K+ in plants). The reports of Bigard et al. (2018; 2020), Duchène et al. (2020), that detailed the genetic diversity for anions (i.e. organic acids) and cations and the consequence in grape acidity in a set of extreme *V. vinifera* varieties and offsprings, here we analyzed the determinants of the acidity of 6 varieties, including 3 sugarless genotypes. As shown in previous sections, sugarless genotypes tend to display a malic acid/tartaric acid ratio higher than the 3 traditional cultivars but with similar sums of malic + tartaric acids and K^+^ levels. As the results, the sugarless genotypes presented similar levels of acidity at the same physiological ripe stage than other varieties.

#### 3. Other cations (Mg^2+^, Ca^2+^)

Magnesium (Mg^2+^) was much less accumulated than K+ in all genotypes. [Mg^2+^] displayed very few changes during ripening (**Table 1, Fig. S6, Fig. S7**). At the arrest of phloem unloading, [Mg^2+^] ranged from 1.6 ^+^/- 1.0 (G5) to 3.7 ^+^/- 1.0 (Merlot N) mEq.L^-1^. Ranging from 4.5 ^+^/- 2.2 (G5) to 5.4 ^+^/- 1.6 (Merlot N) mEq.L^-1^, Calcium (Ca^2+^) was found more accumulated in the green berries than Mg^2+^ (**Table 1**). Then during ripening, [Ca^2+^] tended to decrease (**Fig. S8** and **S9**), ending with concentrations of 1.9 ^+^/- 0.5 (G14) to 3.7 ^+^/- 0.8 (Merlot N) mEq.L^-1^, a relative decrease quite comparable to that of tartaric acid. Therefore, its total amount per berry remains constant during ripening, as widely accepted in grapevine, and consistent with its almost exclusive transport by xylem (Glad et al., 1992; Creasy et al., 1993). Both cations didn’t have a major impact on the wine quality and presented very few variations within the panel of varieties.

**FIg. S6.**
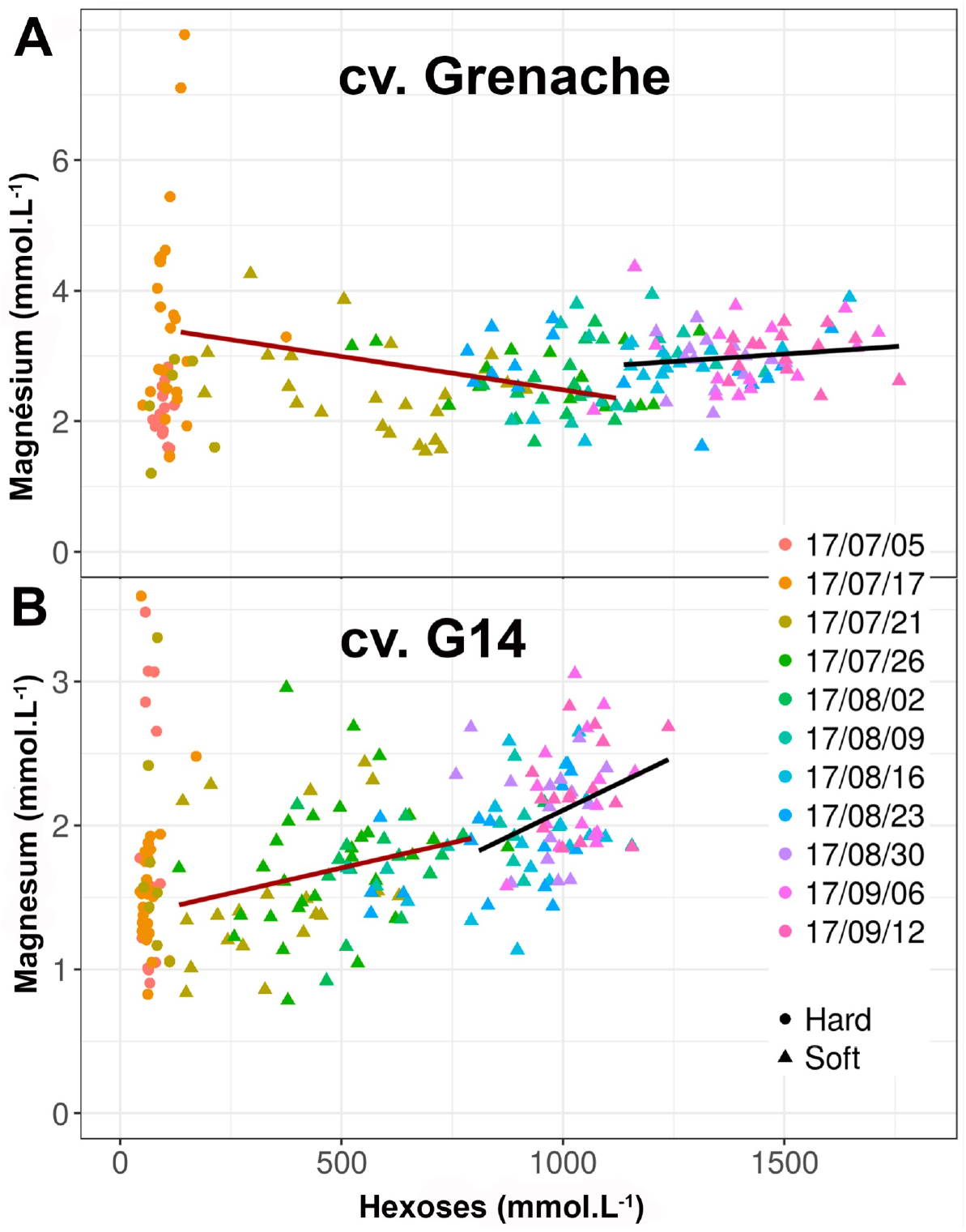
Evolution of the magnesium concentration during berry ripening of Grenache (A) and G14 (B). Lines corresponding to linear fitting during and after phloem unloading.

**FIg. S7.**
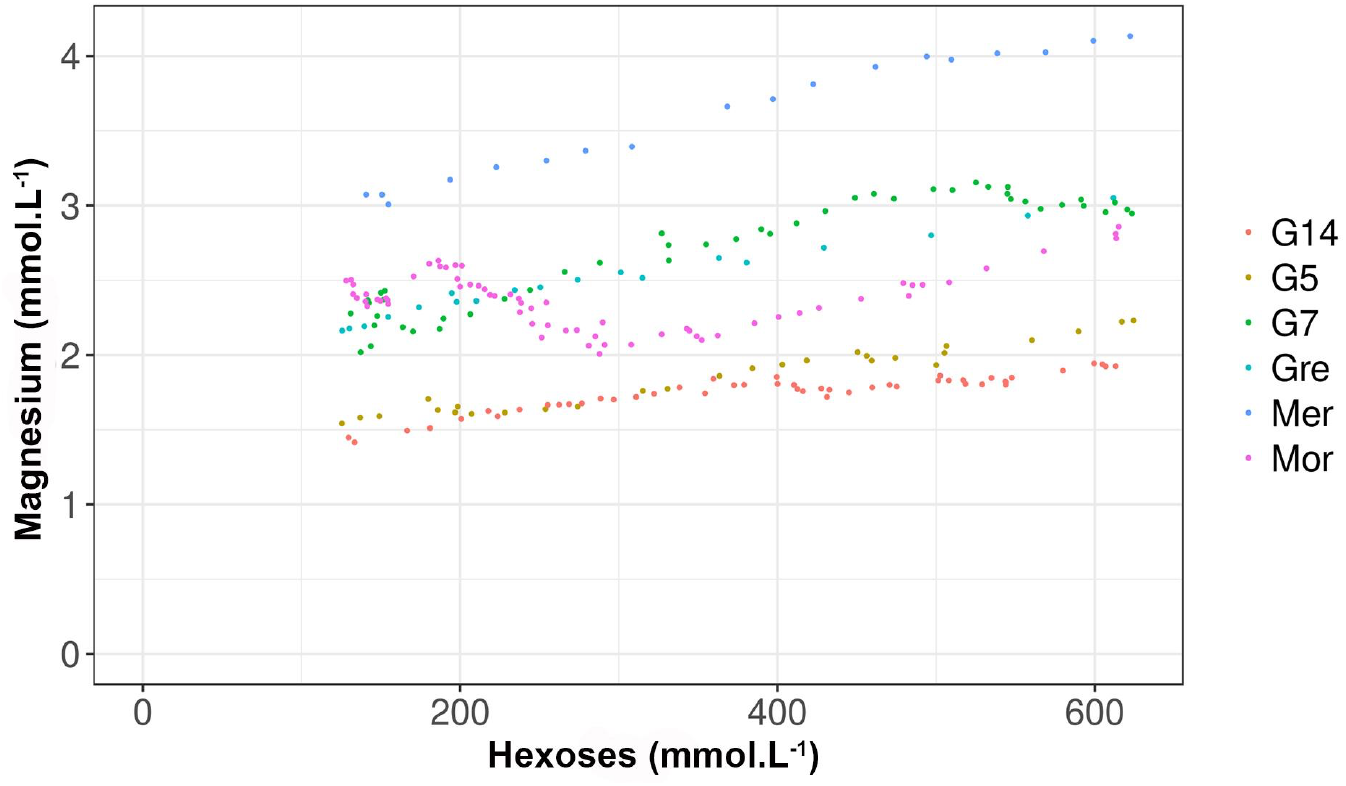
Evolution of the magnesium concentration during early ripening (125-625 mmol.L^-1^ [Hex]) of 6 grapevine varieties.

**FIg. S8.**
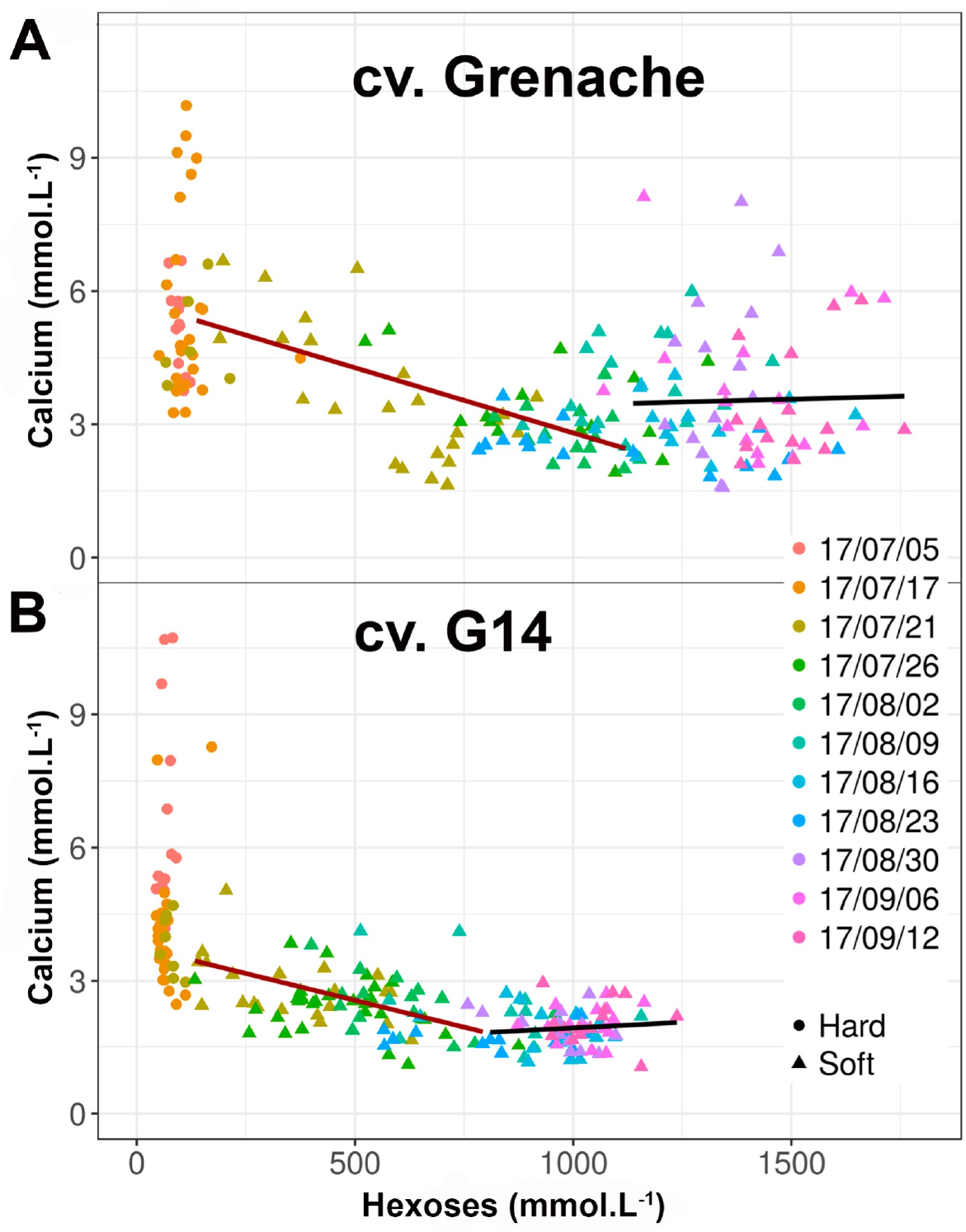
Evolution of the calcium concentration during berry ripening of Grenache (A) and G14 (B). Lines corresponding to linear fitting during and after phloem unloading.

**FIg. S9.**
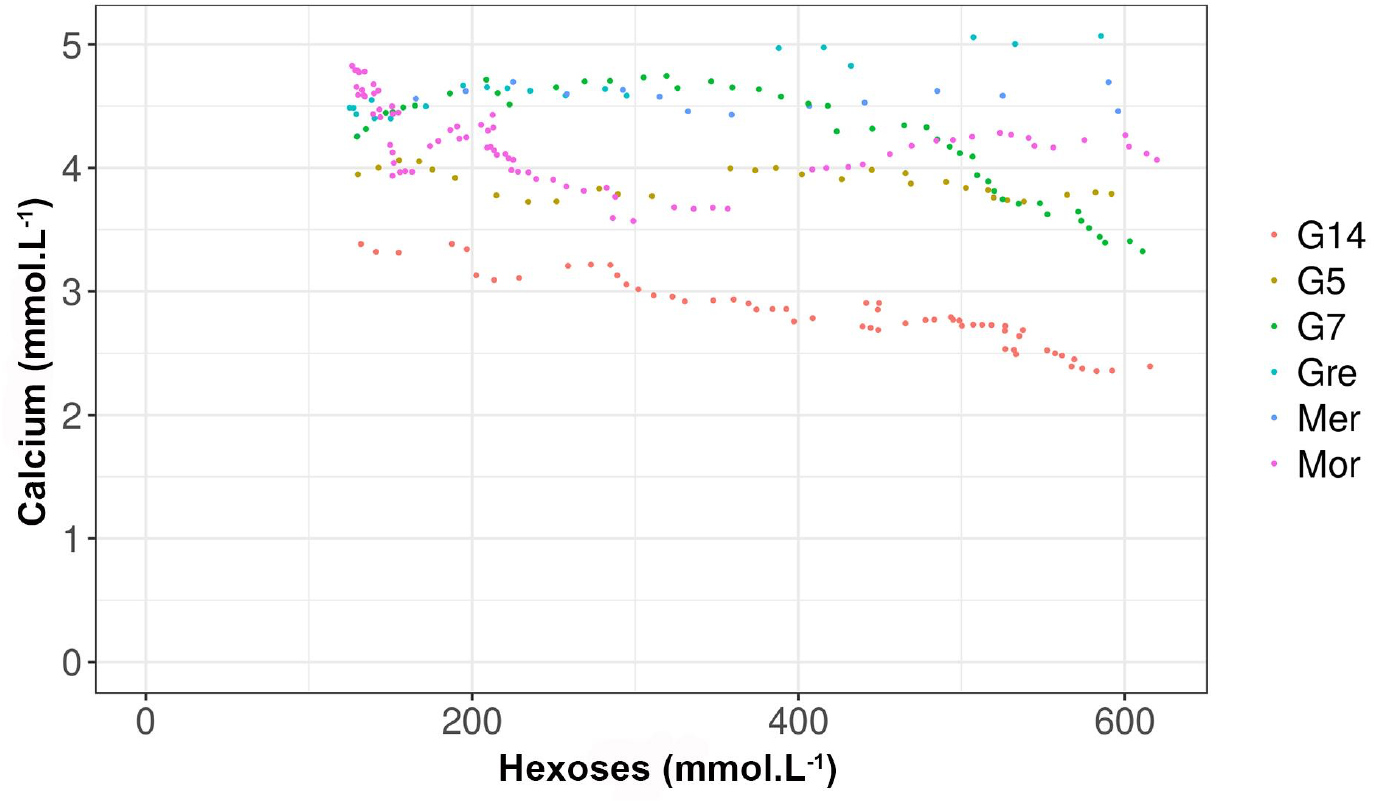
Evolution of the calcium concentration during early ripening (125-625 mmol.L^-1^ [Hex]) of 6 grapevine varieties.

## Conclusion

Getting fruits with reduced [Hex] while preserving their acidity represents an option to mitigate the effect of climate warming on grapevine fruit quality (Torregrosa et al., 2017). This objective can’t be fully addressed by viticultural practices, e.g. harvesting before complete sugar unloading or removing a fraction of the leaves to reduce C photoassimilation, without impacting the quality of the wines (Bobeica et al., 2015; Antalick et al., 2021). Some diversity can be found in *V. vinifera* varieties, or can be obtained by crossbreeding, for water, sugars and the determinants of acidity of the grape (Bigard et al., 2018, 2020). In this study, using advanced methods of berry phenotyping, we have characterized the fruit development and ripening of a set of new disease-tolerant varieties producing low alcoholic wines (Escudier et al., 2017). In previous studies, we have shown that combining single fruit phenotyping and precise physiological landmarks significantly improve the understanding of berry development features (Shahood et al., 2020; Savoi et al., 2021). Indeed, relations between the major solutes are no longer biased upon averaging unsynchronized hence developmentally and metabolically chimeric samples. To circumvent the imprecision of berry growth curves resulting from the heterogeneity in berry size, tartaric acid dilution was used to detect the timing of phloem unloading arrest. This study showed that the sugarless genotypes display a [Hex] reduced by 20-30% when reaching ripe stage without impacting berry growth, organic acid and cations accumulation levels. No major difference being found for fruit growth rates and the quantity of sugars per berry in comparison to control varieties, this suggests the sugarless phenotypes undergo a greater cellular expansion at similar osmotic or turgor pressure. This property is not specific to genotypes deriving from *Muscadinia rotundifolia* and table grape varieties, because Morrastel N also displayed a limited [Hex] in the ripe fruit (< 1 mol.L^-1^). Moreover, similar behaviors can be found in other traditional varieties, such as Aramon, Cornifesto and Mandilaria (Bigard et al., 2018) and Glera, a variety used for Prosecco wine production (https://plantgrape.plantnet-project.org/fr/cepage/Glera). Taken together our results show that adaptive traits to climate changes can be pyramidized with QTLs of tolerance to diseases. By crossing G5 and G14, we have generated microvine segregating progenies (Torregrosa et al., 2019) to further characterize the physiology of this trait and investigate the genetic determinism of water, sugar and organic accumulations (Savoi et al., 2021).

## Acknowledgements

Thanks to Eleoneora Maoddi, Yannick Sire and Philippe Abbal for their contribution in the experiments.

## References

Abbal, P., Boulet, J. C. & Moutounet, M. (1992). Utilisation de paramètres physiques pour la caractérisation de la véraison des baies de raisin. J. Int. Sci. Vigne Vin, 26, 231–237.

Alem, H., Ojeda, H., Rigou, P., Schneider, R. & Torregrosa L. (2021). The reduction of plant sink/source does not systematically improve the metabolic composition of the Vitis vinifera white fruit. Food Chem., 345, 128825. https://doi.org/10.1016/j.foodchem.2020.128825.

Alessandrini, M., Gaiotti, F., Belfiore, N., Matarese, F., D’Onofrio, C. & Tomasi D. (2018). Influence of vineyard altitude on Glera grape ripening (Vitis vinifera L.): effects on aroma evolution and wine sensory profile. J Sci Food Agric., 97, 2695–2705.

Amerine, M. A. & Thoukis, G. (1958). The glucose/fructose ratio of California grapes. Vitis, 1, 224–229.

Amerine, M. A., Roessler, E. B. & Ough, C. S. (1965). Acids and the acid taste. I. The effect of pH and titratable acidity. Am. J. Enol. Vitic., 16, 29–37.

Antalick, G., Šuklje, K., Blackman, J. W., Schmidtke, L. M. & Deloire A. (2021). Performing sequential harvests based on berry sugar accumulation (mg/berry) to obtain specific wine sensory profiles. OenoOne, 55, 131– 146. https://doi.org/10.20870/oeno-one.2021.55.2.4527.

Arrizabalaga, M., Morales, F., Oyarzun, M., Delrot, S., Gomès, E., Irigoyen, J. J., Hilbert, G. & Pascual I. (2918). Tempranillo clones differ in the response of berry sugar and anthocyanin accumulation to elevated temperature. Plant Sci., 267, 74–83. doi: 10.1016/j.plantsci.2017.11.009.

Asproudi, A., Petrozziello, M., Cavalletto, S. & Guidoni S. (2016). Grape aroma precursors in cv. Nebbiolo is affected by vine microclimate. Food Chem., 211, 947–956.

Becard, V., Lacroix, R., Puech, C. & Inaki Garcia de Cortozar-Atauri (2022). Assessment of changes in Grenache grapevine maturity in a Mediterranean context over the last half-century. OenoOne, 56. https://doi.org/10.20870/oeno-one.2022.56.1.4727.

Bigard, A., Berhe, D. T., Maoddi, E., Sire, Y., Boursiquot, J. M., Ojeda, H., Péros, J. P., Doligez, A., Romieu, C. & Torregrosa L. (2018). Vitis vinifera L. fruit diversity to breed varieties anticipating climate changes. Frontiers Plant Sci., doi: 10.3389/fpls.2018.00455.

Bigard, A., Romieu, C., Sire, Y., Veyret, M., Ojeda, H. & Torregrosa L. (2019). Grape ripening revisited through berry density sorting. OenoOne 4, 719–724. DOI:10.20870/oeno-one.2019.53.4.2224.

Bigard, A., Romieu, C., Sire, Y. & Torregrosa L. (2020). Vitis vinifera L. diversity for cations and acidity is suitable for breeding fruits coping with climate warming. Frontiers Plant Sci., DOI: 10.3389/fpls.2020.01175.

Bobeica, N., Poni, S., Hilbert, G., Renaud, C., Gomès, E., Delrot, S. & Dai Z. (2015). Differential responses of sugar, organic acids and anthocyanins tosource-sink modulation in Cabernet Sauvignon and Sangiovese grapevines. Front. Plant Sci., 382. doi:10.3389/fpls.2015.00382.

Burbidge, C. A., Ford, C. M., Melino, V.J., Wong, D.C. J., Jia, Y., Jenkins, C. L. D., Soole, K. L., Castellarin, S. D., Darriet, P., Rienth, M., Bonghi, C., Walker, R. P., Famiani, F. & Sweetman C. (2021). Biosynthesis and cellular functions of tartaric acid in grapevines. Frontiers Plant Sci., 12. doi:10.3389/fpls.2021.643024.

Butrose, M. S., Hale, C. R. & Kliewer W. M. (1971). Effect of temperature on the composition of Cabernet-Sauvignon berries. Am. J. Enol. Vitic., 22, 71–75.

Castellarin, S. D., Gambetta, G. A., Wada H., Krasnow M. N., Cramer G. R., Peterlunger E., Shackel K. A.& Matthews M.A. (2015). Characterization of major ripening events during softening in grape: turgor, sugar accumulation, abscisic acid metabolism, colour development, and their relationship with growth. J. Exp. Bot. 67, 709–722. doi: 10.1093/jxb/erv483.

Champagnol F. (1984). Eléments de physiologie de la vigne et de viticulture générale, ed. Dehan (Montpellier, France: Dehan).

Conde C., Silva P., Fontes N., Dias A., Tavares R.M., Sousa M.J., Agasse A., Delrot S., Gerós H. (2007) Biochemical Changes throughout Grape Berry Development and Fruit and Wine Quality. Food 1: 1–22.

Coombe, B. G. (1976). The development of fleshy fruits. An. Rev. Plant Physiol., 27, 507–528.

Coombe, B. G. (1992). Research on development and ripening of the grape berry. Am. J. Enol. Vitic., 43, 101–110.

Coombe, B. G. & McCarthy, M.G. (2000). Dynamics of grape berry growth and physiology of ripening. Aust. J. Grape Wine Res., 6, 131–135. doi:10.1111/j.1755-0238.2000.tb00171.x.

Cuellar, T., Azeem, F., Andrianteranagna, M., Pascaud, F., Verdeil, J. L., Sentenac, H., Zimmermann, S. & Gaillard I. (2013). Potassium transport in developing fleshy fruits: the grapevine inward K+ channel VvK1.2 is activated by CIPK–CBL complexes and induced in ripening berry flesh cells. Plant J.,73, 1006–1018. doi: 10.1111/tpj.12092.

Dai, Z. W., Ollat, N., Gomès, E., Decroocq, S., Tandonnet, J. P., Bordenave, L. et al. (2011). Ecophysiological, genetic, and molecular causes of variation in grape berry weight and composition: a review. Am. J. Enol. Vitic., 62, 413–425. doi: 10.5344/ajev.2011.10116.

Doumouya, S., Lahaye, M., Maury, C. & Siret R. (2014) Physical and physiological heterogeneity within the grape bunch: impact on mechanical properties during maturation. Am. J. Vitic. Enol., 65, 170–178. doi: 10.5344/ajev.2014.13062.

Duchêne, E., Dumas, V., Jaegli, N. & Merdinoglu D. (2012). Deciphering the ability of different grapevine genotypes to accumulate sugar in berries. Aust. J. Grape Wine Res., 18, 319–328. doi: 10.1111/j.1755-0238.2012.00194.x.

Duchêne, E., Dumas, V., Butterin, G., Jaegli, N., Rustenholtz, C., Chauveau, A., Berard, A., Le Paslier, M. C., Gaillard, I. & Merdinoglu D. (2020). Genetic variations of acidity in grape berries are controlled by the interplay between organic acids and potassium. Theor. App. Genet., https://doi.org/10.1007/s00122-019-03524-9.

Escudier, H., Bigard, A., Ojeda, H., Samson, A., Caillé, S., Romieu, C. & Torregrosa L. (2017) De la vigne au vin : des créations variétales adaptées au changement climatique et résistant aux maladies cryptogamiques 1/2 : La résistance variétale. Revue des Oenologues, 44, 16–18.

Famiani, F., Farinelli, D., Palliotti, A., Moscatello, S., Battistelli, A. & Walker R. B. (2014). Is stored malate the quantitatively most important substrate utilised by respiration and ethanolic fermentation in grape berry pericarp during ripening? Plant Physiol. Biochem., 76, 52–57. doi: 10.1016/j.plaphy.2013.12.017.

Friend, A.P., Trought M.C.T. & Creasy G. L. (2009). The influence of seed weight on the development and growth of berries and live green ovaries in Vitis vinifera L. cvs. Pinot noir and Cabernet-Sauvignon. Aust. J. Grape Wine Res., 15, 166–174.

Gouthu, S., O’Neil, S. T., Di, Y., Ansarolia, M., Megraw, M. & Deluc L.G. (2014). A comparative study of ripening among berries of the grape cluster reveals an altered transcriptional programme and enhanced ripening rate in delayed berries. J. Exp. Bot., 65, 5889–5902. doi: 10.1093/jxb/eru329.

Gutiérrez-Gamboa, G., Garde-Cerdán, T., Carrasco-Quiroz, M., Pérez-Álvarez, E.P., Martínez-Gil, A.M., Del Alamo-Sanza, M. & Moreno-Simunovic Y. (2018). Volatile composition of Carignan noir wines from ungrafted and grafted onto País (Vitis vinifera L.) grapevines from ten wine-growing sites in Maule Valley, Chile. J Sci Food Agric., 98, 4268–4278.

Hawker, J. S., Ruffner, H. P. & Walker R.R. (1976). The sucrose content of some Australian grapes. Am. J. Enol. Vitic., 27, 125–129.

Houel, C., Chatbanyong, R., Doligez A. Rienth, M., Foria, S., Luchaire, N., Roux, C., Adivèze, A., Lopez, G., Farnos, M., Pellegrino, A., This, P., Romieu, C. & Torregrosa L. (2015). Identification of stable QTLs for vegetative and reproductive traits in the microvine (Vitis vinifera L.) using the 18K Infinium chip. BMC Plant Biol., 15,205. DOI 10.1186/s12870-015-0588-0.

Kliewer, W. M. (1966). Sugars and Organic Acids of Vitis vinifera. Plant Physiol., 41, 923–931. http://www.jstor.org/stable/4260763.

Kliewer, W. M. (1967). The glucose-fructose ratio of Vitis vinifera grapes. Am. J. Enol. Vitic., 18, 33–41. doi: 10.1016/j.aca.2011.11.043.

Kliewer, W. M. & Lider L.A. (1970). Effect of day temperature and light intensity on growth and composition of Vitis vinifera L. fruits. J. Amer. Soc Hortic. Sci., 95, 766–769.

Lang, A. & Thorpe M. R. (1989). Xylem, phloem and transpiration flows in a grape: application of a technique for measuring the volume of attached fruits to high resolution using Archimedes’ principle. J. Exp. Bot., 40, 1069–1078. doi: 10.1093/jxb/40.10.1069.

Liang, Z., Sang, M., Fan, P., Wu, B., Wang, L., Duan, W. & Li S. (2011). Changes of polyphenols, sugars, and organic acid in 5 Vitis genotypes during berry ripening. J. Food Sci., 76, 1231–1238.

Liu, H., Wu, B., Fan, P., Xu, H. & Li S. (2006). Sugar and acid concentrations in 98 grape cultivars analyzed by principal component analysis. J. Sci. Food Agric., 86, 1526–1536. doi: 10.1002/jsfa.2541.

Liu, H., Wu, B., Fan, P., Xu, H. & Li S. (2007). Inheritance of sugars and acids in berries of grape (Vitis vinifera L.). Euphytica, 153, 99–107. doi: 10.1007/s10681-006-9246-9.

Matthews, M. A., Cheng, G. & Weinbaum S. A. (1987). Changes in water potential and dermal extensibility during grape berry development. J. Amer. Soc. Hort. Sci., 112, 314–319.

Ojeda, H., Bigard, A., Escudier J. L., Samson, A., Caillé, S., Romieu, C. & Torregrosa L. (2017). De la vigne au vin : des créations variétales adaptées au changement climatique et résistant aux maladies cryptogamiques 2/2 : Approche viticole pour des vins de type VDQA. Revue des Œnologues, 44, 22–27.

Ollat, N., Marguerit, E., Lecourieux, F., Destrac-Irvine, A., Barrieu, F., Dai, Z. et al. (2018). Grapevine adaptation to abiotic stresses. Acta Hortic., 1248. doi: 10.17660/ActaHortic.2019.1248.68.

Parker, A. K., de Cortázar-Atauri, I. G., Gény, L., Spring, J. L., Destrac, A., Schultz H., et al. (2020). Temperature-based grapevine sugar ripeness modeling for a wide range of Vitis vinifera L. cultivars. Agric. For. Meteorol., 285, 107902. doi: 10.1016/j.agrformet.2020.107902.

Ramos, M. C. & Martínez de Toda F. (2021). Interannual and spatial variability of grape composition in the Rioja DOC show better resilience of cv. Graciano than cv. Tempranillo under a warming scenario. OenoOne 55, 85–100. https://doi.org/10.20870/oeno-one.2021.55.3.4695.

R Core Team (2017). R: A Language and Environment for Statistical Computing. Vienna: R Foundation for Statistical Computing.

Rienth, M., Torregrosa, L., Luchaire, N., Chatbanyong, R., Lecourieux, D., Kelly, M., et al. (2014). Day and night heat stress trigger different transcriptomic responses in green and ripening grapevine (Vitis vinifera) fruit. BMC Plant Biol., 14,108. doi: 10.1186/1471-2229-14-108.

Rienth, M., Torregrosa, L., Gauthier, S., Ardisson, M., Brillouet J.L. & Romieu C. (2016). Temperature desynchronizes sugar and organic acid metabolism in ripening grapevine fruits and remodels its transcriptome. BMC Plant Biol., 16, 164. DOI 10.1186/s12870-016-0850-0.

Robin, J. P., Abbal, P. & Salmon J.M. (1997) Firmness and grape berry maturation: definition of different rheological parameters during the ripening. J. Int. Sci. Vigne Vin, 31, 127–138. https://doi.org/10.20870/oeno-one.1997.31.3.1083.

Rogiers, S. Y., Coetzee, Z. A., Walker, R. R., Deloire, A. & Tyerman S.D. (2017). Potassium in the grape (Vitis vinifera L.) berry: transport and function. Frontiers Plant Sci., 8, 1629. https://doi:10.3389/fpls.2017.01629.

Rolle, L., Giacosa, S., Gerbi, V., Bertolino, M. & Novello V. (2013). Varietal comparison of the chemical, physical, and mechanical properties of five colored table grapes. Int. J. Food Prop., 16, 598–612. http://dx.doi.org/10.1080/10942912.2011.558231.

Rösti, J., Schumann, M., Cleroux, M., Lorenzini, F., Zufferey, V., Rienth, M. (2018). Effect of drying on tartaric acid and malic acid in Shiraz and Merlot berries. Aust. J. Grape Wine Res., 24, 421–429. doi: 10.1111/ajgw.12344.

Ruffner, H. P. (1982). Metabolism of tartaric and malic acids in Vitis: A review - Part A. Vitis, 21, 247–259. doi: 10.5073/VITIS.1982.21.247-259.

Santillán, D., Iglesias, A., La Jeunesse, I., Garrote, L. & Sotes V. (2019). Vineyards in transition: A global assessment of the adaptation needs of grape producing regions under climate change. Science of The Total Environment, 657, 839–852. https://doi.org/10.1016/j.scitotenv.2018.12.079.

Savoi, S., Torregrosa, L. & Romieu C. (2021). Transcripts repressed at the stop of phloem unloading highlight the energy efficiency of sugar import in the ripening V. vinifera fruit. Hort. Res., 8, 193. https://doi.org/10.1038/s41438-021-00628-6.

Seguin, B., Stevez, L., Herbin, C. & Rochard J. (2004). Changements climatiques et perspectives pour la viticulture: conséquences potentielles d’une modification du climat. Revue des Oenologues, 111, 59–60.

Shahood, R., Rienth, M., Torregrosa, L. & Romieu C. (2015). Evolution of grapevine (Vitis vinifera L.) berry heterogeneity during ripening. 19th International Meeting of Viticulture GIESCO. Vol. 2, pp. 564–567.

Shahood R. (2017). The berry within an asynchronous harvest: A new paradigm towards the quantitative interpretation of sugar and acid fluxes as major osmoticums and respiratory substrates during bimodal grape development. PhD of Montpellier SupAgro, http://www.theses.fr/s206075, 215 p.

Shahood, R., Torregrosa, L., Savoi, S. & Romieu C. (2020). Berry development hidden by its non-synchronous population. OenoOne, 54, 1077–1092. https://doi.org/10.20870/oeno-one.2020.54.4.3787.

Shiraishi, M., Fujishima, H. & Chijiwa H. (2010). Evaluation of table grape genetic resources for sugar, organic acid, and amino acid composition of berries. Euphytica, 174, 1–13. doi: 10.1007/s10681-009-0084-4.

Storey, R. (1987). Potassium localization in the grape berry pericarp by energy-dispersive x-ray microanalysis. Am. J. Enol. Vitic., 38, 301–309.

Suter, B., Destrac-Irvine, A., Gowdy, M., Dai, Z. & van Leeuwen C. (2021) Adapting wine grape ripening to global change requires a multi-trait approach. Front. Plant Sci., 12, 624867. doi: 10.3389/fpls.2021.624867.

Sweetman, C., Sadras, V. O., Hancock, R. D., Soole, K. L., Ford, C. M. (2014). Metabolic effects of elevated temperature on organic acid degradation in ripening Vitis vinifera fruit. J. Exp. Bot., 65, 5975–5988. doi: 10.1093/jxb/eru343.

Terrier, N., Sauvage, F. X., Ageorges, A. & Romieu C. (2001). Changes in acidity and in proton transport at the tonoplast of grape berries during development. Planta, 213, 20–28. http://www.jstor.org/stable/23386210.

Torregrosa, L., Bigard, A., Doligez A. Lecourieux, D. Rienth, M., Luchaire, N., Pieri, P. Chatbanyong, R., Shahood, et al. (2017). Developmental, molecular and genetic studies on the grapevine response to temperature open breeding strategies for adaptation to warming. OenoOne, 51, 155–165. DOI: 10.20870/oeno-one.2016.0.0.1587.

Torregrosa, L., Rienth, M., Romieu, C. & Pellegrino A. (2019). The microvigne, a model for grapevine physiology studies and genetics. OenoOne, 53. DOI: https://doi.org/10.20870/oeno-one.2019.53.3.2409.

Varandas, S., Teixeira, M. J., Marques, J. C., Aguilar, A., Alves, A., & Bastos M. (2004). Glucose and fructose levels on grape skin: interference in Lobesia botrana behaviour. Analytica Chimica Acta, 513, 351–355., doi:10.1016/j.aca.2003.11.086.

Villete, J., Cuélar, T., Verdeil, J. L., Delrot, S. & Gaillard I. (2020). Grapevine potassium nutrition and fruit quality in the context of climate change. Front. Plant Sci., 11, 123. doi: 10.3389/fpls.2020.00123.

Vondras, A. M., Gouthu, S., Schmidt, J. A., Petersen, A. R. & Deluc, L. G. (2016). The contribution of flowering time and seed content to uneven ripening initiation among fruits within Vitis vinifera L. cv. Pinot noir clusters. Planta, 243, 1191–1202.

Zhang, C., Jia, H., Wu, W., Wang, X., Fang, J. & Wang C. (2015). Functional conservation analysis and expression modes of grape anthocyanin synthesis genes responsive to low temperature stress. Gene, 574, 168–177.

